# LRRK2 modifies α-syn pathology and spread in mouse models and human neurons

**DOI:** 10.1101/522086

**Authors:** Gregor Bieri, Michel Brahic, Luc Bousset, Julien Couthouis, Nicholas Kramer, Rosanna Ma, Lisa Nakayama, Marie Monbureau, Erwin Defensor, Birgitt Schuele, Mehrdad Shamloo, Ronald Melki, Aaron D. Gitler

## Abstract

Progressive aggregation of the protein alpha-synuclein (α-syn) and loss of dopaminergic neurons in the substantia nigra pars compacta (SNpc) are key histopathological hallmarks of Parkinson’s disease (PD). Accruing evidence suggests that α-syn pathology can propagate through neuronal circuits in the brain, contributing to the progressive nature of the disease. Thus, it is therapeutically pertinent to identify modifiers of α-syn transmission and aggregation as potential targets to slow down disease progression. A growing number of genetic mutations and risk factors have been identified in studies of familial and sporadic forms of PD. However, how these genes affect α-syn aggregation and pathological transmission, and whether they can be targeted for therapeutic interventions, remains unclear. We performed a targeted genetic screen of risk genes associated with PD and parkinsonism for modifiers of α-syn aggregation, using an α-syn preformed-fibril (PFF) induction assay. We found that decreased expression of Lrrk2 and Gba modulated α-syn aggregation in mouse primary neurons. Conversely, α-syn aggregation increased in primary neurons from mice expressing the PD-linked LRRK2 G2019S mutation. *In vivo*, using LRRK2 G2019S transgenic mice, we observed acceleration of α-syn aggregation and degeneration of dopaminergic neurons in the SNpc, exacerbated degeneration-associated neuroinflammation and behavioral deficits. To validate our findings in a human context, we established a novel human α-syn transmission model using induced pluripotent stem cell (iPS)-derived neurons (iNs), where human α-syn PFFs triggered aggregation of endogenous α-syn in a time-dependent manner. In PD subject-derived iNs, the G2019S mutation enhanced α-syn aggregation, whereas loss of LRRK2 decreased aggregation. Collectively, these findings establish a strong interaction between the PD risk gene LRRK2 and α-syn transmission across mouse and human models. Since clinical trials of LRRK2 inhibitors in PD are currently underway, our findings raise the possibility that these may be effective in PD broadly, beyond cases caused by LRRK2 mutations.

## Introduction

Parkinson’s disease (PD) is characterized by a progressive loss of midbrain dopaminergic neurons in the substantia nigra pars compacta (SNpc), resulting in motor symptoms, including rigidity, postural instability, tremor at rest and bradykinesia [26, 34]. A defining pathological feature of PD is the presence of intracellular protein aggregates termed Lewy bodies (LB), whose major component is the protein alpha-synuclein (α-syn) [18, 19, 25, 34]. To date, a growing number of dominant and recessive genetic mutations, as well as genetic risk factors, have been identified in families with a high prevalence of PD (fPD), accounting for approximately 5-10% of all PD cases. Further, genome-wide association studies (GWAS) have identified additional genomic loci and candidate genes conferring risk for PD [11]. Rare mutations in *SNCA*, the gene encoding α-syn, as well as duplications and triplications of the locus haven been linked to PD - establishing a direct genetic link between PD and α-syn [4, 12, 35, 36, 50, 54, 64].

α-Syn exhibits certain ‘prion-like’ properties, including the ability to spread through the brain and trigger aggregation of endogenous α-syn in connected neuronal networks [33]. It has been demonstrated that in human PD brains, α-syn pathology is present in the brainstem and olfactory bulb prior to the onset of motor dysfunction, progressively extending to interconnected brain areas such as the SNpc and eventually the neocortex. Importantly, staging of the progressive pathology in human brains correlates with motor symptoms in PD subjects [6, 8, 9]. A variety of *in vitro* and *in vivo* models have been developed, illuminating the transmissive properties of α-syn pathology. In particular, α-syn preformed-fibril (PFF) based rodent models have emerged as a powerful tool to investigate uptake, transport, aggregation and neurodegeneration [10, 21, 40, 60]. Indeed, local injection of α-syn fibrils into the brains of wild type (WT) mice leads to aggregation of endogenous α-syn within neurons of connected regions distant from the injection site [40]. Together, this emerging body of work indicates that α-syn can be taken up by neurons and transmitted to other neurons in interconnected brain areas, where it can trigger aggregation.

Although considerable progress has been made in support of the progressive nature of α-syn pathology, the mechanism underlying the transmission of α-syn aggregates between neurons - and the effect of PD-causing mutations on this process - remains poorly understood [7]. We wondered whether some of the genes that contribute to PD do so by converging on α-syn aggregation and transmission [1]. We performed a series of α-syn PFF-based experiments investigating the effect of a selection of PD-associated risk genes on transmission and aggregation of α-syn pathology in mouse primary neurons, a PFF-based mouse model of PD, and human induced pluripotent stem cell (iPS)-derived neurons from PD patients. In a targeted knock down (KD) and overexpression screen, LRRK2 and GBA, two of the most common genetic risk factors for PD, modulated the extent of α-syn aggregation in mouse primary neurons [31]. *In vivo*, we used an intra-striatal PFF delivery mouse model of PD and observed that α-syn aggregation was exacerbated in transgenic mice carrying the LRRK2 G2019S variant, the most common genetic cause of fPD. In addition to accelerated degeneration of tyrosine hydroxylase (TH)-positive neurons in the SNpc, LRRK2 G2019S transgenic mice also developed more pronounced neuroinflammation and behavioral deficits. To translate our findings in a human context, we employed an approach to rapidly differentiate human iPS cells into induced neurons (iNs) [66] and developed a novel human *in vitro* model of PFF-induced α-syn aggregation. Recombinant human PFFs were rapidly internalized in human iNs and induced robust aggregation of endogenous α-syn. Furthermore, using iPS-derived neurons from fPD patients carrying the LRRK2 G2019S mutation, we observed an increase in α-syn aggregation in human iNs. Conversely, α-syn aggregation was reduced in isogenic LRRK2 knock out (KO) iNs. Collectively, these findings validate our aggregation-based targeted screening approach and identify a strong genetic interaction between LRRK2 and α-syn in both mouse and human PFF-based models of transmission and aggregation. These results have potential implications for clinical trials of LRRK2 inhibitors that are currently underway.

## Materials and Methods

### Animals

Mouse husbandry and procedures were performed in accordance with institutional guidelines and approved by the Stanford Administrative Panel on Animal Care (APLAC). Heterozygous human LRRK2 G2019S BAC transgenic mice (The Jackson Laboratory, cat# 018785) were maintained on a C57BL/6 background (The Jackson Laboratory, cat# 000664). LRRK2 G2019S mice and wild-type litter-mates were used for *in vivo* experiments. C57BL/6 mice, LRRK2 G2019S BAC mice and *CAG–Cas9* transgenic mice (Jackson Laboratory cat# 024858) were used for time-mating and primary neuron cultures.

### Stereotaxic injections

Stereotaxic injections were performed on 8-12 week old adult mice. Animals were placed in a stereotaxic frame and anesthetized with 2% isoflurane (2L/min oxygen flow rate) delivered through an anesthesia nose cone. Ophthalmic eye ointment was applied to prevent desiccation of the cornea during surgery. The area around the incision was trimmed and disinfected. PFF or vehicle solutions were injected unilaterally or bilaterally into the dorsal striatum using the following coordinates (from bregma): anterior = +0.4mm, lateral = +/-1.85mm from midline, depth = −2.7mm (from dura). Mice were injected with sonicated PFFs (5μg/mouse) or PBS vehicle control. PFFs were sonicated prior to injection. Per hemisphere, 1μl volume was injected at a rate of 100nl/min using a 5μl Hamilton syringe with a 32G needle. To limit reflux along the injection track, the needle was maintained in situ for five minutes, before being slowly retrieved. Each mouse was injected subcutaneously with analgesics and monitored during recovery.

### Behavioral testing

Animals underwent a series of behavioral testing up to 6 months post-injection. All behavioral assays were conducted by an investigator blinded to the genotype and treatment condition. Mice were habituated to the behavioral testing room and investigator prior to the behavioral testing. All testing equipment was cleaned with Virkon and 70% ethanol between animals.

-*Rotarod* was used to assess motor learning and coordination. To acclimate them to the Rotarod apparatus, mice were trained on 3 consecutive days for 5min at constant speed of 10rpm. For the testing, thee speed was gradually increased from 4-40rmp over the course of 5min. Latency to fall was measured for 3 trials/day on 3 consecutive days. The rod and chambers were wiped with 70% ethanol between trials.

-*Pole test* was used to measure motor coordination and function. Mice were placed on a vertical pole and the latency to descend was measured for 3 trials.

-*Wire hang test* was used to test motor function and deficit. Mice were placed on a grid that was then inverted. The latency to fall was recorded for two consecutive trials.

-*Open Field* was used to measure spontaneous activity, locomotion and anxiety. Animals are placed in the square open field arena (76cm x 76cm x 50cm) and allowed to freely move while being recorded by the automated tracking system (Ethovision) for up to 10 minutes. Distance traveled, number of entries and time spent in the center and periphery was automatically assessed.

### Tissue processing

Mice were anesthetized with isoflurane and transcardially perfused with 0.9% saline. Brains were dissected and fixed in 4% paraformaldehyde (PFA) pH 7.4, at 4°C for 48 hours. Brains were then stored in 30% sucrose in 1x PBS at 4°C. For unilateral injection experiments, the whole brain was fixed in PFA. For bilateral injection experiments, one hemibrain was fixed in PFA whereas the other hemibrain was sub-dissected, snap frozen in liquid nitrogen and stored at −80°C for biochemical analysis. PFA fixed brains were sectioned at 35μm (coronal sections) with a cryo-microtome (Leica) and stored in cryoprotective medium (30% glycerol, 30% ethylene glycol) at −20°C.

### Immunohistochemistry

Tissue processing and immunohistochemistry was performed on free-floating sections according to standard published techniques. 1:4 to 1:12 series coronal sections were used for all histological experiments. Sections were rinsed 3 times in TBST, pre-treated with 0.6% H2O2 and 0.1% Triton X-100 (Sigma-Aldrich) and blocked in 5% goat serum in TBST. The following primary antibodies were used: NeuN (1:1000, Millipore, Cat# MAB377), Tyrosine Hydroxylase (TH; 1:1000; Pel-Freez, cat# P40101-0), α-syn pSer129 (81A; 1:5000, Covance/BioLegend cat# MMS-5091); α-syn pSer129 (1:3000; Abcam cat# 51253), LRRK2 (1:1000; NeuroMAP cat# 75-253), p62 (1:1000; Proteintech cat# 18420-AP), GFAP (1:1000; Dako cat# Z0334), Iba1 (1:1000; Wako cat# 019-19741), CD68 (1:500; AbD Serotec cat# MCA1957GA), C1q (1:1000; Abcam cat# ab182451). After overnight incubation at 4°C, the primary antibody staining was revealed using biotinylated secondary antibodies, ABC kit (Vector Laboratories) and 3,3’-diaminobenzidine tetrahydrochloride (DAB, Sigma-Aldrich). Fluorescently labeled secondary antibodies were used for fluorescent and confocal microscopy. Sections were mounted on Superfrost Plus slides (Fisher Scientific) and coverslipped using Entellan (for DAB staining) or ProlongDiamond antifiade mountant (for fluorescent immunostaining). Imaging and unbiased stereological estimations of TH and pSer129-positive cells were performed by investigators blinded to genotype and experimental conditions. Per animal 8-10 SNpc sections were used for TH-neuron analysis. 3-5 sections per animal and brain region were used for all other immunostainings.

### Thioflavin S staining

Sections were rinsed 3 times in PBS, mounted on Superfrost Plus slides and dried overnight. The slides with the brain sections were then incubated in 0.1% Thioflavin S solution in 20% Ethanol for 30 minutes at room temperature. The samples were rinsed twice with 20% Ethanol for two minutes, followed by 2 washes in water. Following the Tioflavin S staining, sections were labeled for pSer129 using standard immunohistochemistry techniques and mounted using Vectashield without DAPI (Vector Labs cat# H-1000). Fluorescence images were captured using a ZEISS LSM 800.

### Western blotting

Mouse brain tissue was subdissected, snap frozen and stored at −80°C until further use. *In vitro*, iPS cells and neurons were washes with ice-cold PBS prior to lysis. Cells or tissue were lysed on ice in 1x RIPA lysis buffer or sequentially in 1% Triton X followed by 2% SDS lysis buffer supplemented with 1x Halt protease and phosphatase Inhibitor Cocktail (Thermo Fisher Scientific cat# 78429, 78426). [59]. Crude RIPA lysates were centrifuged at 15,000 *g* for 10 min at 4°C to remove cellular debris. TritonX-lysates were centrifuged at approximately 15,000g for 10min at 4°C. The pellet was resuspended and sonicated in 2% SDS lysis buffer. Clarified lysates were quantified with a Pierce BCA protein assay (Thermo Fisher Scientific cat# 23225). Cell lysates were mixed with 4x NuPage LDS loading buffer or 4x Laemmli buffer and equal amounts of protein were subjected to SDS–PAGE, transferred to PVDF or nitrocellulose membranes. For the detection of total endogenously expressed α-syn (Novus Biologicals cat# NBP-92694) and α-syn pSer129 (Abcam cat# 51253), PVDF membranes were fixed with 0.4% PFA for 30 minutes at room temperature prior to the blocking step [37]. Membranes were blocked with 5% milk or 5% BSA in Tris-Buffered Saline with 0.1% Tween (TBST) and immunoblotted according to standard protocols. The following additional antibodies were used: ACTB (clone C4; 1:5000; Millipore cat#), GAPDH (1:5,000, Sigma cat# G8795), total soluble α-synuclein (1:1000, Novus Biologicals cat# NBP-92694), α-synuclein pSer129 (Abcam cat# 51253, 1:5000), LRRK2 (1:1000, Abcam cat# ab133474), GBA (1:1000; Novus Biologicals cat# NBP1-32271), Synaptophysin 1 (SYSY cat# 101 002), PSD95 (1:5000; Abcam cat# ab18258), Homer 1 (SYSY cat# 160 003). Following incubation at 4°C overnight, horseradish peroxidase-conjugated secondary antibodies and an ECL kit (GE Healthcare/Amersham Pharmacia Biotech) were used to detect protein signals. Multiple exposures were taken to select images within the dynamic range of the film (GE Healthcare Amersham Hyperfilm ECL). Protein bands were quantified using FIJI software (NIH). Actin or GAPDH bands were used for normalization.

### RT-qPCR

RNA was isolated from brain tissue and cell pellets using TRIZol reagent (Thermo Fisher Scientific, Cat# 15596026) and PureLink™ RNA Mini Kit following the manufacturer’s instructions. The RNA concentration was determined via Nanodrop and RNA was reverse transcribed using the High-Capacity cDNA Reverse Transcription Kit (Thermo Fisher Scientific, Cat# 4368813). Real time PCR was performed on a Applied Biosystems StepOnePlus Real-Time PCR instrument using 2x TaqMan Universal Master Mix (Cat # 4440040) and gene-specific TaqMan probes against Snca (mouse, Mm01188700_m1), Lrrk2 (mouse, Mm00481934_m1), ActB (Mm02619580_g1), Ifna1 (mouse, Mm03030145_gH), Ifnb1 (mouse, Mm00439552_s1), C3 (mouse, Mm01232779_m1), C1qa (mouse, Mm00432142_m1), Il6 (mouse, Mm00446190_m1), Il1b (mouse, Mm00434228_m1), Tgfb1 (mouse, Mm01178820_m1), Il10 (mouse, Mm01288386_m1), Tnf (mouse, Mm00443258_m1), Trem2 (mouse, Mm04209424_g1), Vim (mouse, Mm01333430_m1), Cxcl10 (mouse, Mm00445235_m1), Cd44 (mouse, Mm01277161_m1), Gfap (mouse, Mm01253033_m1), SNCA (human, Hs01103383_m1), LRRK2 (human, Hs01115057_m1), GRIA1 (human, Hs00181348_m1), MAPT (human, Hs00902194_m1), MAP2 (human, Hs00258900_m1), SNAP25 (human, Hs00938957_m1). ActB was used for normalization. Each sample and primer set was run in triplicates and relative expression level were calculated using the ΔΔCT method.

### PFF preparation

The expression and purification of mouse or human wild-type α-syn was performed as previously described [24]. α-Syn fibril formation was induced by incubation in 50mM Tris–HCl, pH 7.5, 150mM KCl buffer at 37°C under continuous shaking in an Eppendorf Thermomixer at 600rpm. α-Syn fibrils were centrifuged twice at 15,000g for 10min and resuspended in PBS. For *in vitro* uptake and transport experiments, the fibrils were labeled with Alexa-488 or Alexa-555 (Life Technology, # A-20009) NHS fluorophore following the manufacturer’s instructions using a protein:dye ratio of 1:2. The labeling reactions were arrested by addition of 1mM Tris pH 7.5. Free fluorophores were removed by two cycle or centrifugations at 15,000g for 10min and resuspension of the pelleted fibrils in PBS. All fibrils were fragmented prior to *in vivo* or *in vitro* use by sonication for 20 min in 2-ml Eppendorf tubes in a Vial Tweeter powered by an ultrasonic processor UIS250v (250 W, 2.4 kHz; Hielscher Ultrasonic, Teltow, Germany). For human α-syn PFFs, 6.5μg/ml (0.5μM) were used for aggregation experiments and 13μg/ml (1μM) were used for internalization experiments with human induced neurons. For mouse α-syn PFFs, 5ug of fibrils/mouse were used for *in vivo* mouse experiments. 2.6μg/ml (0.2μM) were used for aggregation experiments and 6.5μg/ml (0.5μM) were used for internalization experiments in primary mouse neurons.

### Virus production

HEK293T (ATCC) cells were cultured under standard conditions (DMEM, 10% FBS, penicillin–streptomycin) and used to package lentiviral particles according to standard protocols with third-generation packaging plasmids (pMDlg-pRRE, pRSV-ReV, pMD2.0; Addgene). Lentiviral backbone and packaging plasmids were transfected into HEK293T cells using Lipofectamine 2000 and Lentivirus-containing medium was harvested after 48h. Media was centrifuged at 300g for 5min to remove cellular debris, and concentrated using ultracentrifugation (26,000rpm/2hrs/4°C) or Lenti-X concentrator (Clontech) before being added to cell cultures. ShRNAs were expressed from H1 promoters (pGreenPuro backbone, SystemBio), gRNAs were expressed from U6 promoters. For overexpression experiments, ORFs from the human ORFeome collection were cloned into the pLEX307 destination plasmid (Addgene cat# 41392) using the standard LR clonase protocol (Thermo Fischer Scientific).

### Primary Neuron cultures

Primary mouse hippocampal and cortical neurons were dissociated into single-cell suspensions from E16.5 mouse brains with a papain dissociation system (Worthington, Cat# LK003153). Neurons were seeded onto poly-L-lysine–coated plates (0.1% (wt/vol)) and grown in Neurobasal medium (Gibco) supplemented with B-27 serum-free supplement (Gibco), GlutaMAX, and penicillin–streptomycin (Gibco) in a humidified incubator at 37°C, with 5% CO2. Half media changes were performed every 4-5d, or as required. Neurons were plated on 12mm glass coverslips (Carolina Biological Supplies cat# 633009) in 24-well plates (100,000 cells/well), fin 12-well plates for biochemical studies (500,000/well), or in 96-well plates for automated imaging and luminescence based survival assays (25,000-35,000 cells/well). For the targeted shRNA-based screens, neurons were grown in clear bottom, black wall plates (Sigma cat# CLS3340-50EA), transduced on 1-3 day in vitro (DIV). For gene-knockout experiments in Cas9 cortical neurons, lentiviruses encoding sgRNAs expressed from a U6 promoter were transduced on 1DIV. Sonicated mouse PFFs were added to neurons on 10DIV. 2.6μg/ml PFFs were used for shRNA-based screens and 6.5μg/ml for internalization experiments. Neurons were fixed or lysed on 17-21DIV.

### ES/iPS and iN culture conditions

ES and iPS cells were cultured using feeder-free conditions on Matrigel (Fisher Scientific cat# CB-40230) using mTeSR1 media (Stemcell Technologies, cat# 85850). ROCK inhibitor Y-27632 was added to the culture media during passaging. Cells were transduced with a Tet-On inducible system allowing for the expression of the transcription factor Ngn2 upon addition of doxycycline to the media. Cells were differentiated as previously described in DMEM/F12 media supplemented with 1x N2 (ThermoFisherScientific) and 1ug/ml doxycycline [66]. Cells were dissociated on day 3-5 of differentiation and replated on poly-L-lysin coated tissue culture plates, glass coverslips or microfluidics chambers (Xona microfluidics SND450). iNs were transitioned to Neurobasal media with 1x B27 and doxycycline starting on day 6. Half media changes were performed every 3-4d, or as required.

### Internalization of labelled alpha-synuclein fibrils

The internalization experiments were performed as previously described [10]. Primary mouse neurons or induced neurons were grown in 8 well Lab Tek II chamber slides (ThermoFisherScientific cat# 155409). For induced human neurons, sonicated Alexa-488 labeled α-synuclein fibrils were added at a final concentration of 13μg/ml (1μM) after 14-21 days of differentiation. For primary neurons labeled mouse fibrils were added at a final concentration of 6.5μg/ml (0.5 μM). Internalization was measured at 1, 2, 4, 6, 24 h after the addition of fibrils to the culture media. At each time point, the medium was removed, the cells were washed gently 2-3 times with PBS. 0.1% Trypan blue (ThermoFischerScientific, T10282) was added to each well to quench the non-internalized fluorescent signal and the cells were imaged using a Leica DMI6000B inverted fluorescence microscope with a 20x objective. Control wells without fibrils were used to measure the background fluorescence. 20 images were acquired per well and the percentage of neurons containing fluorescent puncta as well as the average intensity per neuron was measured using ImageJ. More than 100 neurons were assessed for each well.

### Immunocytochemistry

Cells were grown on poly-l-lysine–coated glass coverslips (0.1% (wt/vol) in standard multi-well cell culture plates and were stained through standard immunocytochemistry techniques. Briefly, cells were fixed with 4% formaldehyde for 15 minutes at room temperature, rinsed with PBS, permeabilized with 0.1% Triton X-100, blocked with 5% normal goat serum in PBS. The following primary antibodies were used for labeling: NeuN (1:1000, Millipore cat# MAB377), α-syn pSer129 (1:5000; Covance cat# MMS-5091), α-syn pSer129 (1:3000, Abcam ab51253), α-syn (1:1000, Abcam cat# ab138501), p62 (1:1000; Proteintech cat# 18420-AP), ubiquitin (1:500, Millipore cat# MAB1510), Map2 (1:1000; SYSY cat# 188 004), Map2A/B (1:1000; Millipore cat# MAB378), B-III-tubulin (1:1000; SYSY cat# 303 304). After over-night primary antibody incubation, cells were rinsed 3x with PBS. Cells were incubated with fluorescently labeled secondary antibodies for 1h at room temperature. Coverslips were mounted with Prolong Diamond Antifade mountant with DAPI (Thermo Fisher Scientific). Images were acquired with a Leica DMI6000B inverted fluorescence microscope and a confocal Zeiss LSM710 microscope. Automated imaging and analysis were performed with the IncuCyte life cell analysis imaging system and Molecular Devices ImageXpress Micro and MetaXpress 5. The percentage area of MAP2, BIIItubulin and pSer129 immunofluorescence staining was quantified in ImageJ or MetaXpress 5. The number of NeuN and DAPI labeled-cells were counted using MetaXpress 5 and the analyze particle function in FIJI. The experimentors were blinded to the treatment conditions. For the targeted shRNA based screen, neurons were grown in clear bottom, black wall plates optimized for automated imaging. Images were acquired in a 4×3 grid pattern for each well (12-24 wells/shRNA/screen). The area covered by pSer129 and the number of NeuN-positive nuclei were measured for each image using MetaXpress. Scramble shRNA and untreated control wells were included on each plate. The pSer129 aggregation score was calculated using the pSer129 intensity and area and normalized to the NeuN counts. Scramble controls were included in all plates and used as comparison between plates.

### Statistics

Statistical tests were performed with GraphPad Prism 7. Differences between treatment conditions were established using a unpaired Student’s t test (for two conditions). For experiments with > 2 groups, a one-way ANOVA with a Tukey’s post test for multiple comparisons or two-way ANOVA with Bonferroni correction was performed. Pearson r was calculated for correlation analysis. p < 0.05 was considered statistically significant. Statistic details are indicated in the respective figure legends.

## Results

### A targeted genetic screen in mouse primary neurons identifies interaction between familial PD genes and α-syn aggregation

Overexpression of WT α-syn in mouse primary neurons was well tolerated and did not lead to acute degenerative phenotypes (Fig. S1a-c). Moreover, transmission of α-syn could be investigated in the absence of overt degeneration using recombinant α-syn PFFs (Fig. S1a,d,e). In this *in vitro* model, the addition of PFFs to neurons induces aggregation of endogenously expressed α-syn, with synaptic changes and degenerative phenotypes observed only at late stages [42, 60]. We exposed cortical mouse neurons to sonicated α-syn PFFs (Fig. S2) at 10 days *in vitro* (DIV) and detected aggregation of endogenously expressed α-syn using phospho-Serine129-specific (pSer129) immunolabeling. We first observed pSer129 in neurites after 3 days exposure to PFF. The amount of pSer129 increased progressively over time (Fig. S1d,e). We established an automated imaging-based screening approach that combined detection of pSer129 and the neuronal marker NeuN with targeted manipulation of a selection of PD-associated risk genes (Fig. 1a).

**Fig. 1.**
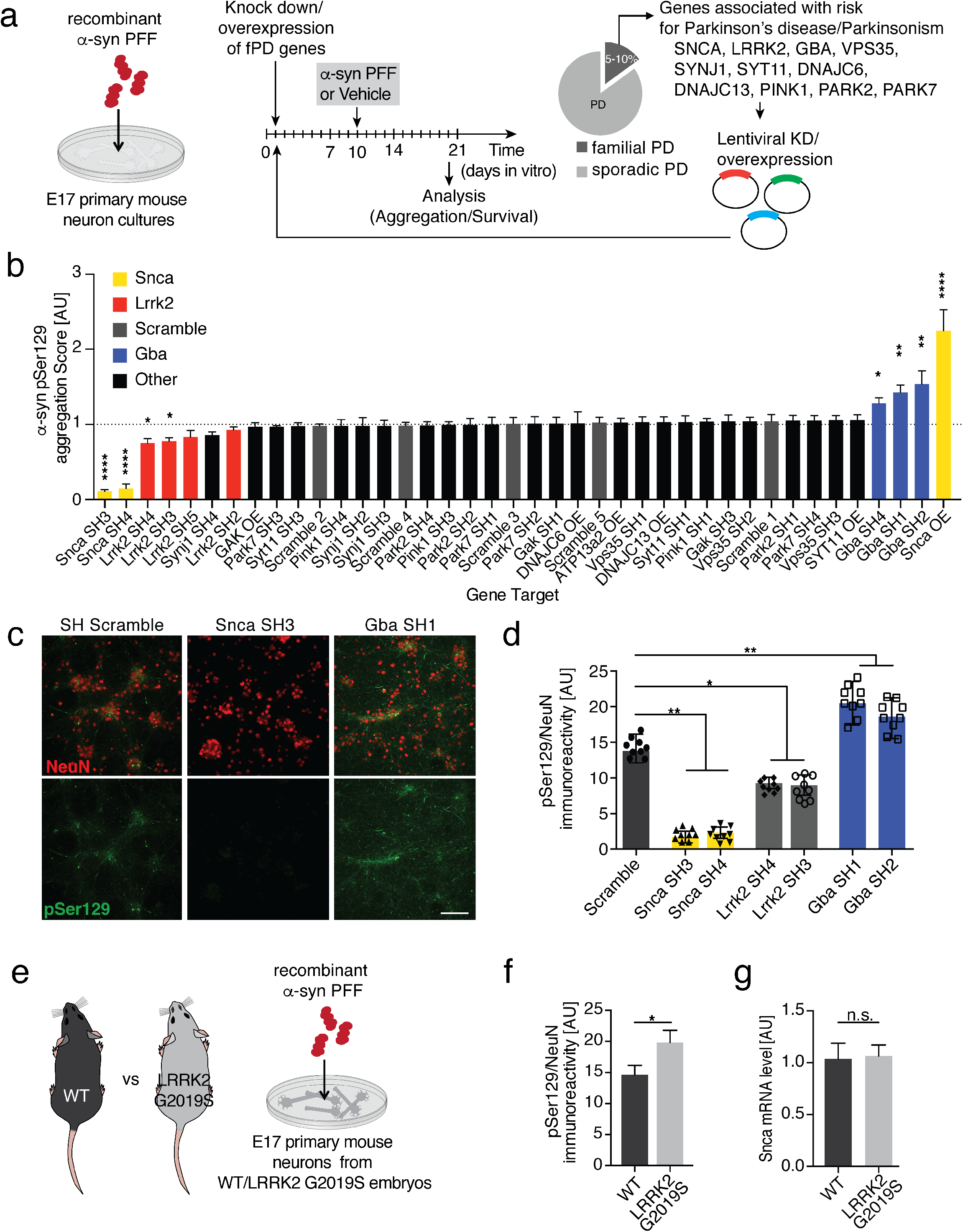
PFF-based screen in primary neuron identifies Lrrk2 and Gba as genetic modifiers of α-synuclein transmission and aggregation. **a** Schematic of experimental design: lentiviral knockdown or overexpression of Parkinson’s disease (PD) risk genes in primary neuron cultures exposed to fragmented α-syn preformed fibrils (PFFs) at 10 days in vitro (DIV). Survival and aggregation of endogenously expressed α-syn was measured using immunolabeling of phosphorylation of α-syn at Serine 129 (pSer129) 10 days after PFF treatment. **b** Summary of targeted screening results. The aggregation score was calculated based on pSer129 immunolabeling and normalized to the neuronal viability for each shRNA. Neuronal viability was determined by the number of NeuN-positive cells/well in PFF compared to vehicle treated wells. The dotted line marks the average of all non-targeting scramble shRNAs. A minimum of 2 shRNAs with a KD-efficiency of 50 percent were used per PD risk gene. α-syn (Snca) constructs are highlighted in yellow, Lrrk2 in red, Scramble controls in gray and Gba in blue (n=12-24 wells/shRNA). Data expressed as mean + SEM; *p < 0.05; **p < 0.01;****p<0.0001 compared by one-way ANOVA with a Tukey’s posttest for multiple comparisons. **c,d** Representative images (**c**) and quantification of pSer129-positive (green) α-syn aggregation (**d**) following knock down of *Snca, Gba, Lrrk2* and nontargeting scramble (SCR) control (n=8-10 independent wells/treatment). pSer129 immunolabeling was normalized to neuron counts using the neuronal marker NeuN (red). **e,f** Experimental design (**e**) and quantification of PFF-based α-syn aggregation (**f**) in primary neuron cultures derived from LRRK2 G2019S-mutant (G2019S) and WT mouse embryos (n=10 wells/genotype, 10 images/well). **g** Relative α-syn (Snca) mRNA expression level measured in WT and G2019S-LRRK2 mutant neurons at 10DIV using quantitative RT-qPCR analysis (n=5 samples/group). Data expressed as mean + SEM; *p < 0.05; **p < 0.01; compared by unpaired Student’s t-test and oneway ANOVA with a Tukey’s post-test for multiple comparisons.

We developed lentiviral shRNA knock down (KD) and overexpression plasmids targeting PD-linked genes, including *Gba1, Lrrk2, Vps35, Synj1, Syt11, Gak, Atp13a2, Pink1, Park2, Park7, Dnajc6, and Dnajc13* (Fig. 1a and Fig. S3). We infected mouse primary neurons at 1 DIV and measured shRNA-mediated KD at 10 DIV. We included shRNAs that achieved at least 50% KD in further screening. Previously, the level of α-syn expression has been shown to determine the extent of aggregation and mediate selective vulnerability of neuronal populations in PFF-based models of α-syn transmission and aggregation [40, 42, 54, 56, 60]. To assess the dynamic range of the screening assay, we included *Snca* KD and overexpression conditions as controls. Consistent with previous reports, *Snca* KD led to a significant reduction in pSer129 levels (Fig. 1b-d) [42, 60], while neuronal overexpression of α-syn led to a two-fold increase (Fig. 1b).

For the screen, we infected primary cortical neurons at 1 DIV, added fragmented α-syn PFF at 10 DIV and assessed neuronal pSer129 staining at 20 DIV (Fig. 1a). Using pSer129 as a read out, we observed that Gba KD significantly increased α-syn aggregation in primary neurons (Fig. 1c,d). Gba KD also led to an increase in total α-syn levels in the absence of PFF treatment, likely accounting for the observed increase in α-syn aggregation (Fig. S4a,b). A potential feedback loop between Gba and α-syn level has been reported in neurons [44]. Interestingly, several PD-associated genes such as *Pink1, Parkin, Synj1*, associated with early onset parkinsonism without real evidence of synucleinopathy, did not impact synuclein aggregation in this screening assay (Fig. 1b). Lastly, *Lrrk2* KD led to a significant decrease in pSer129 without changing the total level of α-syn (Fig. 1b-d and Fig. S4a,b). We further validated the effect of *Lrrk2* KD using Lrrk2-targeting gRNAs in Cas9 transgenic mouse primary neurons. Consistent with our shRNA results, gRNAs targeting endogenous *Lrrk2* significantly reduced pSer129 in primary neurons (Fig. S4c,d). Importantly, *Lrrk2* gRNAs did not alter α-syn levels in the absence of PFF treatment (Fig. S4e). Together, these data identify genetic interactions between several familial PD genes and α-syn aggregation.

### LRRK2 G2019S mutation increases α-syn aggregation in mouse primary neurons

Mutations in LRRK2, in particular the G2019S mutation, are the most common genetic cause of PD [47, 68]. LRRK2 mutations, including G2019S are thought to cause disease by a gain-of-function mechanism, likely via increased kinase activity [27, 62]. Therefore, we tested whether expression of LRRK2 G2019S would increase α-syn aggregation in the PFF model. We used primary cortical neurons derived from human BAC transgenic mice, expressing LRRK2 under the control of the endogenous human regulatory elements (Fig. 1e, Fig. S4f) [61]. Using the PFF-based aggregation assay, we observed a significant increase in pSer129 levels in neurons from LRRK2 G2019S transgenic mice compared to WT controls (Fig. 1f). Importantly, LRRK2 G2019S did not alter expression of endogenous α-syn and other synaptic proteins in primary neurons (Fig. 1g and S4g,h). Additionally, we did not observe any differences in cellular toxicity between LRRK2 G2019S mutant and WT PFF-treated neurons (Fig. S4i,j). Considering that several Rab GTPases have recently been identified as LRRK2 kinase substrates, potentially modulating α-syn uptake, trafficking, escape from the endo-lysosomal compartment and aggregation, we next measured the internalization kinetics of α-syn fibrils [3, 5, 20, 55]. However, we did not observe any significant differences in uptake or endocytosis of α-syn PFFs between LRRK2 G2019S mutant and WT neurons (Fig. S4k). These data indicate that expression of LRRK2 G2019S is associated with increased α-syn aggregation without impacting internalization of PFF or expression of endogenous α-syn in mouse primary neurons.

### PFF injection into the striatum induces α-syn aggregation and degeneration in mice

Previous reports have demonstrated that local PFF delivery into the striatum of rodents induces aggregation of endogenously expressed α-syn in remote brain areas projecting to the injection site, including the SNpc [40, 49]. PFF-based *in vivo* mouse models allow for both the ability to perform longitudinal studies, as well as to assess diverse neuronal populations and the complex interplay between neurons and glial cells in the context of neurodegeneration [42, 63]. We harnessed this model to test the impact of LRRK2 G2019S on α-syn aggregation and degeneration. We first used unilateral stereotaxic injections to deliver fragmented mouse PFFs or vehicle control into the striatum of non-transgenic adult mice, followed by behavioral testing and histological analysis up to 6 months post injection (Fig. S5a). Consistent with previous reports [40], PFF-injected mice developed widespread pSer129-positive, p62-positive and Thioflavin S-positive inclusions in both hemispheres (Fig. S5b-f) in regions that include the cortex, amygdala, and striatum. Importantly, TH-positive dopaminergic neurons in the SNpc, an area associated with movement dysfunction in human PD, developed robust α-syn pathology ipsilateral to the injection site (Fig. S6a-c). Approximately 30% of TH-positive neurons contained pSer129-positive inclusions by 3 months post injection, followed by a decline between 4-6 months post injection (Fig. S6c). Further, we observed a significant decrease in the number of TH-positive neurons in the SNpc 4-6 months post injection, paralleling the decline in TH-positive pSer129-positive double labeled neurons (Fig. S6d-f). In contrast to previous reports [39, 40, 43, 49, 63], we did not observe robust deficits in behavioral paradigms testing motor control and coordination (Fig. S6g). However, mice injected with PFFs displayed altered anxiety-related behavior in the open field test at 6 months post injection. While there was no difference in overall activity and distance travelled, mice injected with PFFs spent significantly more time in the center as opposed to vehicle injected mice (Fig. S6h). Cumulatively, the PFF-based mouse model mimicked key histopathological features of PD.

### LRRK2 G2019S expression exacerbates PFF-mediated α-syn aggregation and degeneration pathology in mice

Having established baselines for the PFF-based *in vivo* model, we next investigated the role of human LRRK2 G2019S in aggregation of α-syn in the basal ganglia circuitry, degeneration of TH neurons and behavior. We used human LRRK2 BAC-transgenic mice, expressing the G2019S mutant form of LRRK2 using human regulatory elements (Fig. 2a,b). LRRK2 G2019S transgenic mice developed normally and no significant differences in α-syn expression, motor function and body weight were detected in up to 12-month-old transgenic animals compared to WT littermates (Fig. S7). For subsequent experiments, LRRK2 G2019S transgenic mice and WT littermates were stereotaxically injected bilaterally into the striatum with α-syn PFFs or vehicle control, followed by behavioral testing and histological analysis at 1, 3 and 6 months post-injection (Fig. 2a,b).

**Fig. 2.**
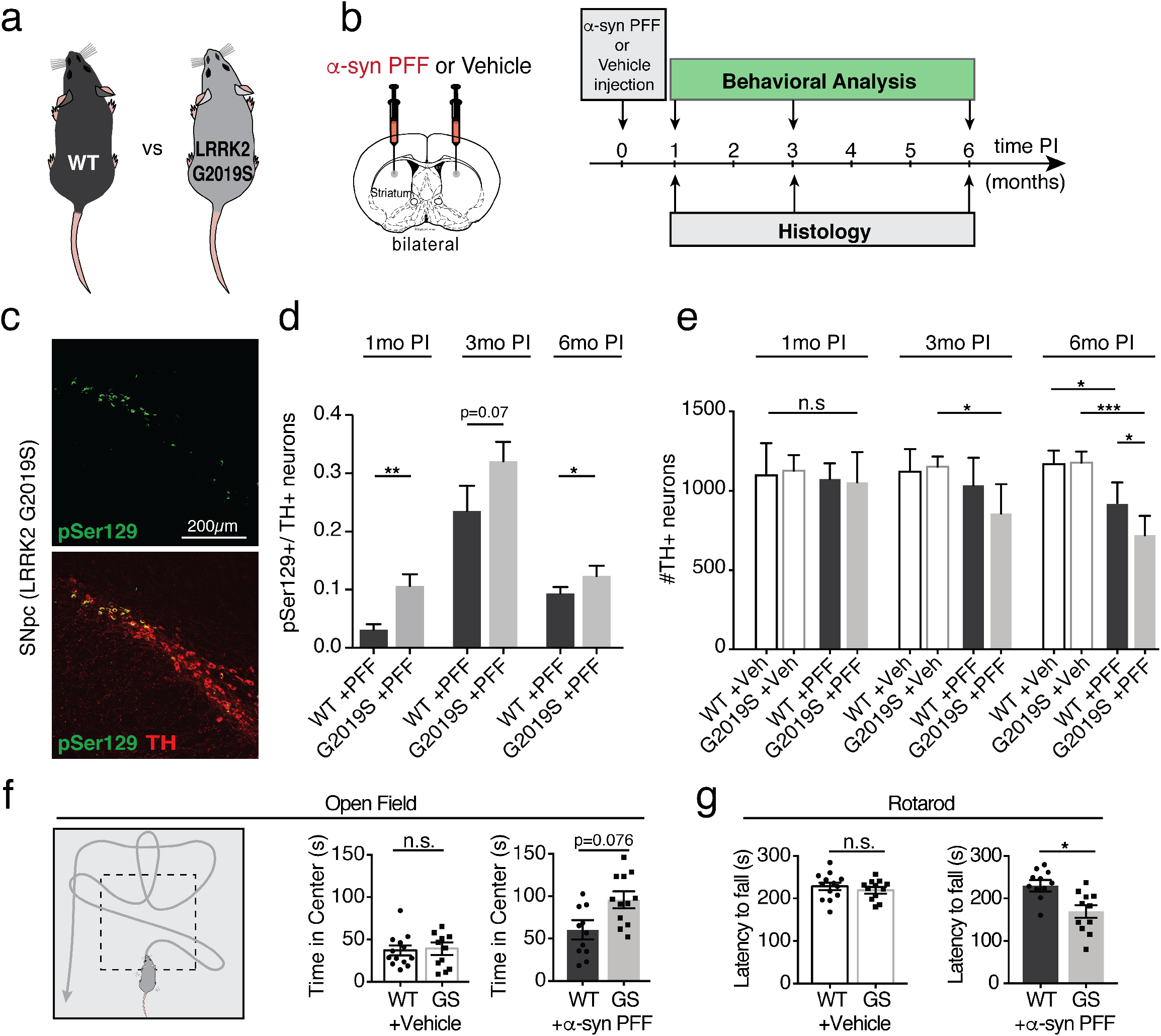
The LRRK2 G2019S mutation exacerbates α-syn pathology in a PFF-based *in vivo* mouse model of α-syn transmission and aggregation. **a,b** Schematic of mouse genotypes and experimental design. Adult LRRK2 G2019S BAC-transgenic mice (G2019S or GS) and wildtype (WT) littermates were injected with α-syn PFFs or vehicle control (Veh). Histological analysis and behavioral testing was performed at 1,3,6 months post injection (PI). **c** Representative images of pSer129 (green) immune-labeling in TH-positive neurons (red) in the substantia nigra pars compacta (SNpc) at 6 months PI. **d** Quantification of the percentage of TH-positive neurons that are also pSer129-positive in the SNpc of PFF-injected animals (n=8-13 animals/group, 8-10 brain sections/animal). **e** Quantification of the number of TH-positive neurons in the SNpc of WT and LRRK2-G2019S animals at 1,3,6 months post injection with PFFs or vehicle control (n=6-12 animals/group, 8-10 brain sections/animal). **f** Schematic and results of open field behavioral testing 6 months post injection with PFFs or vehicle control (11-13 animal/group). **g** Rotarod behavioral assay measuring the latency to fall from an accelerating rotating cylinder at 6 months post injection (n=11-13 animals/group). Data expressed as mean + SEM; *p < 0.05; compared by Student’s t-test and one-way ANOVA with a Tukey’s post-test for multiple comparisons.

At 1 month post-injection, the LRRK2 G2019S transgenic mice displayed significantly more pSer129 aggregates in the dorsal striatum, motor cortex and amygdala. However, by 3 and 6 months post-injection, both LRRK2 G2019S transgenic animals and WT controls developed widespread pSer129-positive inclusions in the brain (Fig. S8). No pSer129-positive inclusions were detected in either vehicle injected LRRK2 G2019S transgenic or WT control animals at any time point (Fig. S8). In the SNpc, the percentage of TH-positive pSer129-positive neurons was significantly higher in LRRK2 G2019S transgenic mice at 1 month post injection compared to WT controls (Fig. 2c,d). Although a significant difference in pSer129 expression persisted up to 6 months post-injection, the relative differences between the two groups decreased over time (Fig. 2c,d). As previously observed in WT animals (Fig. S6c), the percentage of TH-positive dopaminergic neurons co-labeled with pSer129 decreased between 3 and 6 months post-injection for both genotypes (Fig. 2d). Using stereological analysis of TH-positive neuronal cell bodies in the SNpc, we observed accelerated loss of TH-positive neurons in LRRK2 G2019S transgenic mice compared to WT littermate controls at 3 months post-injection that was further exacerbated at 6 months post-injection (Fig. 2e). Importantly, we did not observe any degeneration of TH-neurons in vehicle injected animals (Fig. 2e). Together, these findings indicate an accelerated degenerative phenotype in LRRK2 G2019S transgenic mice, with increased α-syn pathology and loss of TH-positive neurons occurring in the SNpc at early time points compared to WT littermate controls.

Next, we investigated whether increased α-syn aggregation and dopaminergic neuron degeneration observed in PFF-injected LRRK2 G2019S transgenic mice was associated with behavioral deficits. At 6 months post injection, using the open field test we observed that PFF-injected LRRK2 G2019S transgenic mice spent more time in the center than WT control animals (Fig. 2f). Additionally, using rotarod testing, we assessed motor behavior and coordination. Surprisingly, we observed a significant decrease in the ability of LRRK2 G2019S transgenic mice to stay on a rotating cylinder compared to WT controls (Fig. 2g). Collectively, these behavioral data indicate decreased anxiety phenotypes and increased motor impairments in α-syn PFF-injected LRRK2 G2019S transgenic mice.

Lastly, protein aggregation and neuronal cell loss is often associated with neuroinflammation in human neurodegenerative diseases and mouse models of neurodegeneration. Moreover, a recent study demonstrated a role for microglia and astrocytes in the development of α-syn pathology in mouse models of PD [63]. Given ubiquitous expression of LRRK2 in the human and mouse CNS, including in neurons and glial cells [65], we next examined changes in glial activation in LRRK2 G2019S transgenic and WT animals following PFF injection. While we detected increased microglia activation in both PFF-injected groups, we observed altered expression of microglial markers Iba1 and Cd68, between LRRK2 G2019S transgenic mice and WT littermate controls (Fig. 3a-c). Additionally, using RT-qPCR analysis, we observed increased levels of inflammatory markers Il6, Tnfa and C1qa, as well as astrocytic markers Vim, Cd44 and Cxcl10 in LRRK2 G2019S transgenic animals compared to WT controls (Fig. 3d-f). Histologically, we also observed elevated C1q expression in PFF injected LRRK2 G2019S transgenic mice compared to WT controls (Fig. 3g,h). Together, these data identify accelerated degeneration, exacerbated behavioral impairments and accompanying neuroinflammatory phenotypes in PFF-injected LRRK2 G2019S transgenic mice.

**Fig. 3.**
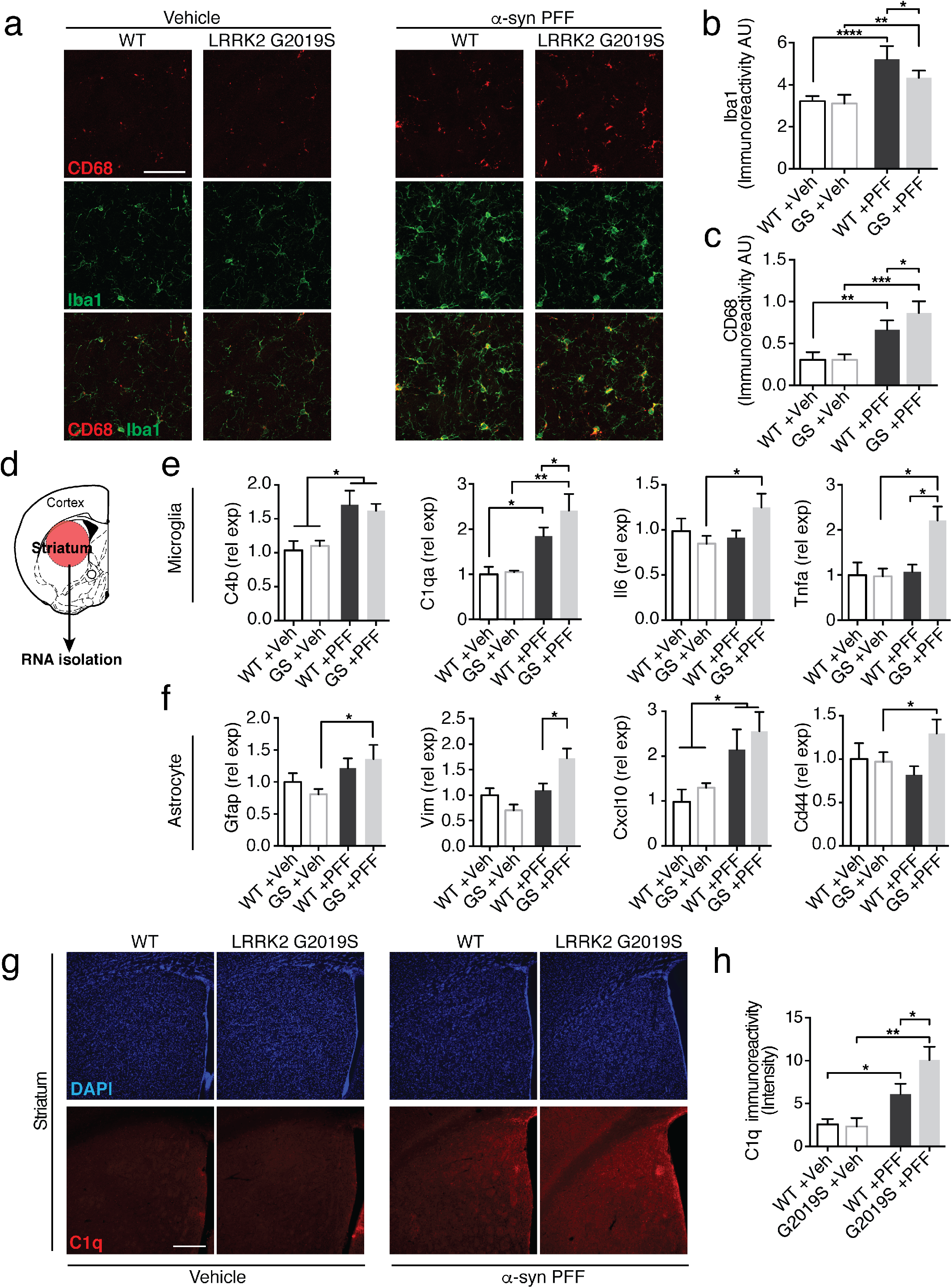
The neuroinflammatory response is altered in PFF-injected LRRK2 G2019S mutant mice. a-c Representative images (**a**) and quantification (**b,c**) of microglial markers Iba1 (green) and Cd68 (red) in the dorsal striatum of LRRK2 G2019S (G2019S or GS) mice and wildtype (WT) littermate controls 6 months post-injection (PI) with PFFs or Vehicle (Veh) control (n=6-12 animals/group). **d-f** Experimental design and quantification of gene expression of microglial (**e**) and astrocyte-associated (**f**) activation markers. mRNA was isolated from the striatum and gene expression was analyzed using RT-qPCR analysis (n=5-6 animals/group). **g-h** Representative images (**g**) and quantification (**h**) of intensity of C1q immunolabeling in the dorsal striatum of PFF and vehicle injected mice 6 months post injection (n=5-6 animals/group). Data expressed as mean + SEM; *p < 0.05, **p < 0.01; compared by Student’s t-test, one-way ANOVA with a Tukey’s post-test for multiple comparisons or and two-way ANOVA with Bonferroni post hoc correction.

### Establishing a model of PFF-transmitted α-syn aggregation in human induced neurons

In addition to rodents, preformed fibril based aggregation models have been reported in marmosets and rhesus macaques, demonstrating that transmission in primate models is possible [41, 48, 51, 53]. However, the translation of these findings to human cells has been limited by the lack of robust and scalable human neuron-based assays. To evaluate the role of LRRK2 in α-syn transmission in the context of human pathology, we developed a novel *in vitro* model of PFF-induced α-syn aggregation in human neurons. A key limitation in PFF-based models is the dependency on expression of endogenous α-syn [40, 42, 56]. Moreover, due to its predominant pre-synaptic localization α-syn expression is further dependent on neuronal maturation and synaptogenesis. Differentiation and maturation of human ES/iPS-derived neurons is often a relatively inefficient, laborious and lengthy process, yielding heterogenous populations of neurons that are not ideal for PFF-based *in vitro* models. To circumvent technical limitations and avoid the confounds linked to α-syn overexpression, we employed an induced neuron (iN) approach previously shown to rapidly differentiate human ES/iPS cells into functional cortical neurons (Fig. S9a) [66]. Human iPS cells differentiated rapidly into iNs with neuronal morphology, expressing neuronal markers NeuN, BIII-tubulin, TAU and MAP2 (Fig. S9b,c). Additionally, α-syn expression significantly increased over the first 2 weeks of differentiation (Fig. S9d,e), in line with neurite and synaptic markers (Fig. S9f-i).

Previous reports indicate a species-dependent barrier between mouse and human fibrils, potentially impacting cross-seeding and templating efficiency [39]. Therefore, we treated human neurons with fragmented human α-syn PFFs. Consistent with our mouse primary neuron experiments, we added human PFFs to 2-3 week-old iNs and performed pSer129 immunolabeling at various time points following fibril treatment (Fig. 4a). Starting at approximately 6-10 days post fibril treatment, we detected elongated and serpentine-like pSer129 staining in PFF - but not vehicle-treated iNs. Levels of pSer129 labeling significantly increased with time (Fig. 4b,c), resembling α-syn inclusions observed at early stages in mouse primary neurons treated with mouse fibrils, although the pathology developed with slower kinetics in human iNs (Fig. S1d,e). Elongated pSer129-positive structures resembled Lewy neurites and predominantly co-localized with the axonal markers BIII-tubulin and Tau (Fig. 4d). Phosphorylated aggregates also partially co-localized with ubiquitin, similar to observations in mouse primary neurons (Fig. 4e) [60]. Additionally, we sequentially extracted and immunoblotted lysates from human iNs exposed for 14 days with human PFFs in 1% Triton X followed by 2% SDS, and detected pSer129 predominantly in the SDS resistant fraction. Collectively, these data indicate that PFFs trigger aggregation of endogenous α-syn in human iNs.

**Fig. 4.**
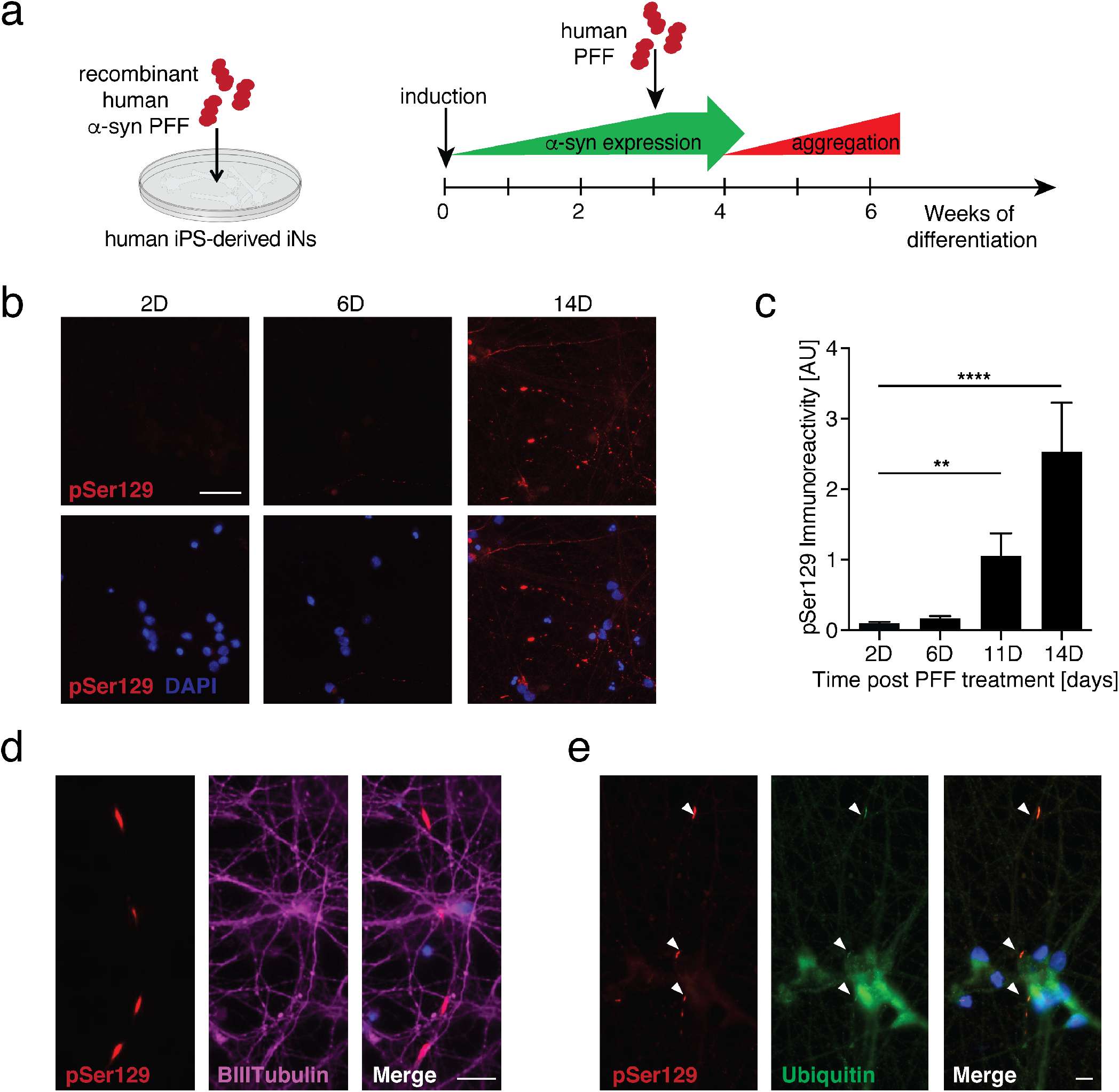
Recombinant human α-syn PFFs induce aggregation of endogenous α-syn in human iPS-derived induced neurons. **a** Experimental design: iPS-derived induced neurons (iNs) were differentiated for 3 weeks before the addition of human α-syn PFFs to the culture media. Aggregation of endogenously expressed α-syn was assessed using immunolabeling of pSer129. **b,c** Representative images (**b**) and quantification (**c**) of pSer129 immunolabeling (red) up to 14 days post PFF treatment (n=19-21 frames/time point). Nuclei labeled with DAPI (blue). **d** Representative images of pSer129 α-syn (red) co-localizing with the axonal marker BIII-tubulin (magenta) 7 days post PFF treatment. **e** Representative images of pSer129 α-syn (red) co-localizing with ubiquitin (green) 14 days post PFF treatment. Data expressed as mean + SEM; compared by one-way ANOVA with a Tukey’s post-test for multiple comparisons.

### LRRK2 G2019S mutation and LRRK2 expression modulate α-syn aggregation in fPD patient-derived human induced neurons

We next used our human neuron-based PFF model to investigate uptake and aggregation of α-syn in LRRK2 mutant iNs. We used fPD patient-derived human iPS cells from LRRK2 G2019S carriers, isogenic gene corrected lines and isogenic LRRK2 KO lines [52]. We first confirmed expression of LRRK2 and the presence of the G2019S mutation in all isogenic lines (Fig. S10a). We differentiated all lines and did not observe any difference in survival between G2019S mutant, corrected and LRRK2 KO human iNs (Fig. S10b,c). We then measured the internalization of α-syn PFFs by human iNs (Fig. 5a-c). After 24 hours, approximately 95% of human iNs contained fluorescently labeled α-syn PFFs. We did not observe any significant difference between LRRK2 mutant and isogenic corrected lines, indicating that LRRK2 expression and activity did not significantly alter α-syn PFF internalization in human iNs (Fig. 5c).

**Fig. 5.**
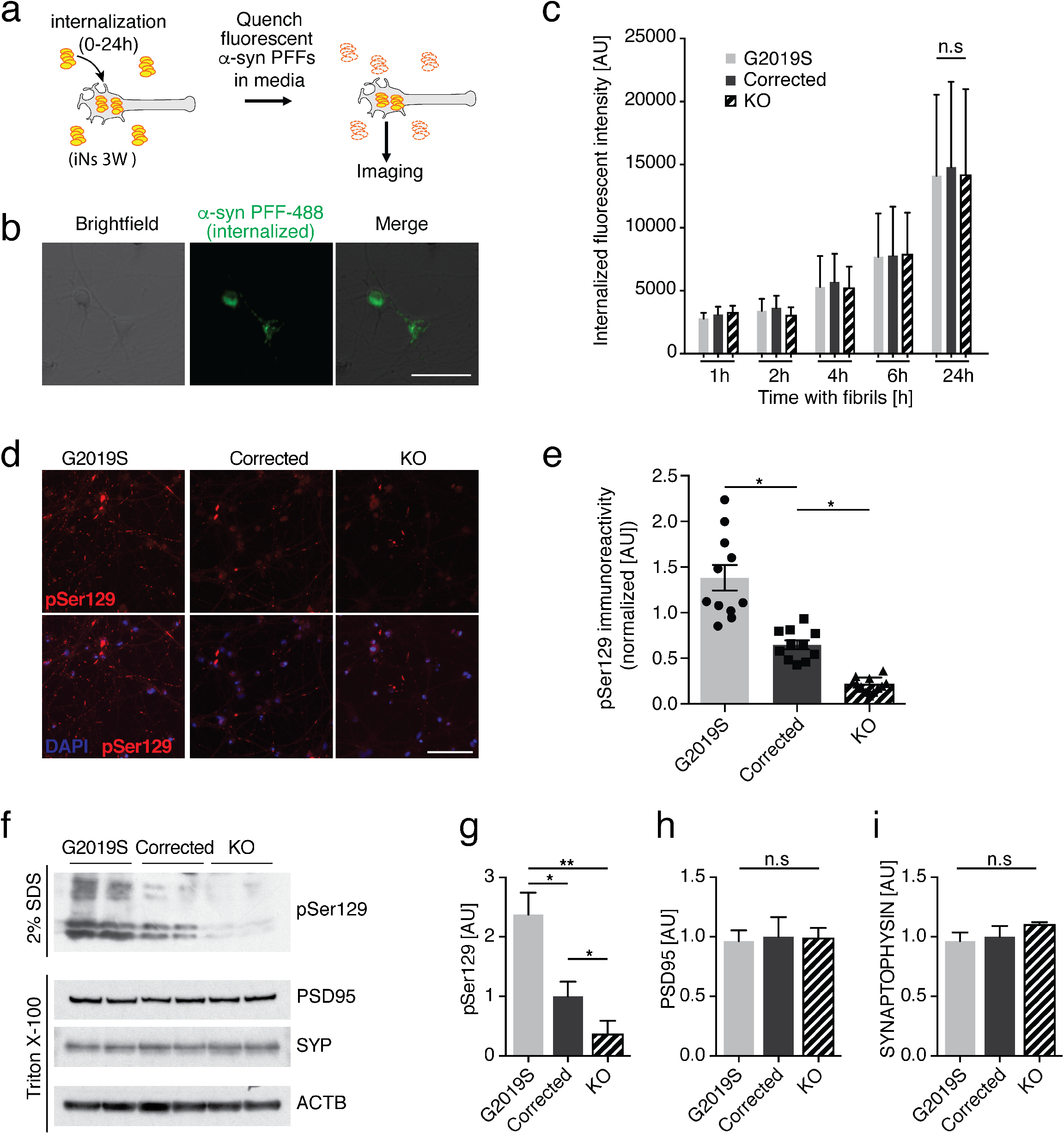
LRRK2 mutations and level modulate aggregation of α-syn in human induced neurons. **a** Experimental design of internalization of α-syn PFFs. 3 week old induced neurons (iNs) were exposed to fluorescently labeled α-syn PFFs for 0 to 24 hours. Fluorescence in the media was quenched with 0.1% Trypan blue, revealing the internalized labeled PFFs. **b** Representative image of internalized PFFs (green) in human iNs (brightfield) after 6 hours of incubation. **c** Quantification of internalized PFFs in isogenic G2019S mutant, corrected and LRRK2 knock out (KO) iNs after 1-24 hours of incubation (n=300-450 cells from two biological replicates/condition). Data expressed as mean + SD, compared by two-way ANOVA with Bonferroni post hoc correction. **d,e** Representative images and quantification of pSer129-positive α-syn aggregates after 3 weeks of incubation with recombinant human PFFs (n=10-12 wells/group). Data expressed as mean + SEM; *p < 0.05; **p < 0.01; compared by one-way ANOVA with a Tukey’s post-test for multiple comparisons. **f** Representative Western blot of isogenic LRRK2 G2019S mutant, corrected and LRRK2 knock out (KO) iN lysates 2 weeks post PFF treatment, probed with anti-pSer129, anti-PSD95, anti-SYNAPTOPHYSIN (SYP) and anti-ACTB antibodies. Synaptic proteins and ACTB were used as loading controls. Proteins were sequentially extracted using 1% Triton-X followed by 2% SDS lysis buffers. **g-i** Quantification of pSer129 (**g**), PSD95 (**h**) and SYNAPTOPHYSIN (**i**) 14 days post PFF treatment (n = 4/group). Data expressed as mean + SEM; *p < 0.05; **p<0.01; compared by one-way ANOVA with a Tukey’s post-test for multiple comparisons.

We next used the PFF transmission model to investigate aggregation of endogenously expressed α-syn. We compared pSer129 labeling between G2019S mutant, corrected and LRRK2 KO human iNs. We observed a significant increase in pSer129 in G2019S mutant human iNs, consistent with our findings in mouse primary neurons (Fig. 5d,e and Fig. 1e,f). Surprisingly, LRRK2 KO human neurons displayed a significant reduction in pSer129 (Fig. 5d,e). We corroborated these findings in lysates from human iNs two weeks post exposure to fibril, detecting increased pSer129 in G2019S mutant compared to corrected human iNs in the 2% SDS fraction. Conversely, pSer129 was decreased in lysates from LRRK2 KO human iNs (Fig. 5f,g). We did not observe any significant differences in the pre- and postsynaptic markers Synaptophysin and PSD95 in PFF treated human iNs (Fig. 5h,i). Given the extent of α-syn aggregation is sensitive to endogenous expression level, we measured and detected comparable α-syn levels between G2019S mutant and corrected human iNs independent of PFF treatment (Fig. 6a,b). Surprisingly, the amount of α-syn was decreased in LRRK2 KO compared to G2019S mutant and corrected human neurons, potentially accounting for decreased pSer129 levels observed in LRRK2 KO iNs. Due to the synaptic localization of endogenous α-syn, we next examined whether there were baseline differences in neuronal maturation and synaptogenesis between G2019S mutant, corrected and LRRK2 KO human iNs. Interestingly, we did not observe any differences in the levels of other synaptic markers, including Synaptophysin, Homer1 and PSD95 (Fig. 6b-e). Furthermore, the differentiation-dependent upregulation of neuronal and synaptic markers was comparable between the different human isogenic lines, as assessed using previously identified pan-neuronal and mature neuron markers (Fig. 6f-i) [57]. Together, these data demonstrate that PFFs can seed aggregation of endogenously expressed α-syn in fPD-derived human iNs. Importantly, both the G2019S mutation, as well as LRRK2 expression level, modulated α-syn aggregation in this PFF-based human iNs *in vitro*, indicating a conserved role for genetic interaction between fPD genes and α-syn aggregation across mouse and human models.

**Fig. 6.**
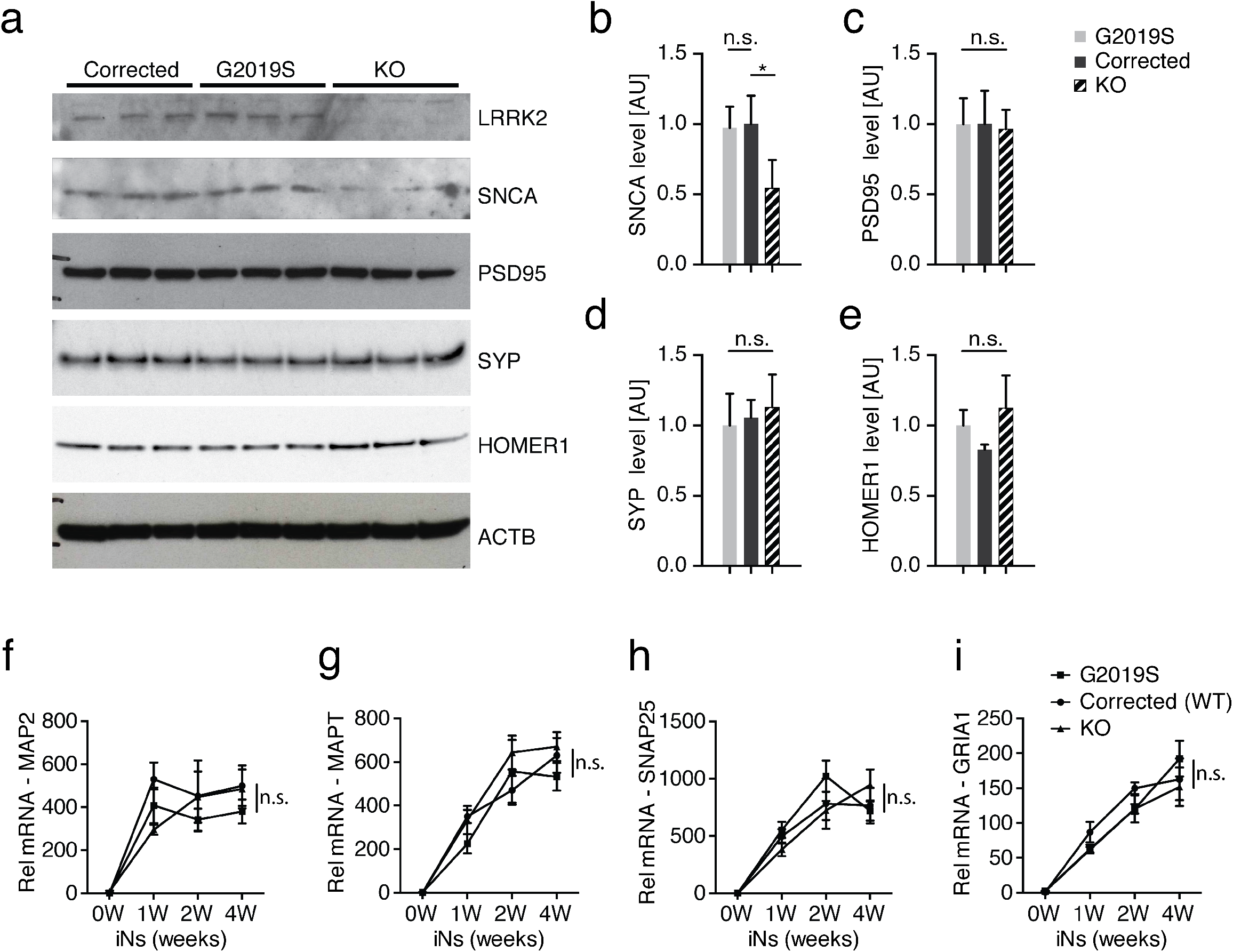
α-syn expression is altered in human LRRK2 knock-out induced neurons. **a** Representative Western blot of lystates from isogenic LRRK2 G2019S mutant, corrected and knock out (KO) induced neurons (iNs) at 4 weeks of differentiation probed with anti-LRRK2, anti-α-syn (SNCA), anti-PSD95, anti-SYNAPTOPHYSIN (SYP), anti-HOMER1 and anti-ACTB antibodies. ACTB was used as a loading control. **b-e** Quantification of α-syn (**f**), PSD-95 (**g**), SYNAPTOPHYSIN (SYP) (**d**) and HOMER1 level at 4 weeks of differentiation (n=3/group). Data expressed as mean + SD; compared by one-way ANOVA with a Tukey’s post-test for multiple comparisons. **f-i** qRT-PCR expression analysis of mature neuron markers MAP2 (**f**), MAPT (**g**) and synaptic markers SNAP25 (**h**) and GRIA1 (**i**) during 4 weeks of differentiation. Data normalized to the 0 weeks of differentiation time point (n=4/group and time point). Data expressed as mean + SEM; *p < 0.05; **p<0.01; compared by one-way ANOVA with a Tukey’s post-test for multiple comparisons and two-way ANOVA with Bonferroni post hoc correction.

## Discussion

In our study, we employed a targeted screen of PD-associated risk genes to identify LRRK2 as a mediator of induced α-syn aggregation. We demonstrated that abrogation of endogenous Lrrk2 in mouse primary neurons decreases α-syn aggregates, whereas expression of the familial PD-associated G2019S mutation increases aggregation. *In vivo*, transgenic mouse data demonstrated that expression of LRRK2 G2019S exacerbates α-syn aggregation, dopaminergic neuron degeneration, as well as associated neuroinflammatory and behavioral impairments in a PFF-based mouse model of PD. We extended these observations by developing and characterizing a novel *in vitro* human neuron PFF-based transmission and aggregation model. Using human iPS-derived induced neurons, we observed that human neurons rapidly internalize recombinant human PFFs, triggering the aggregation of endogenously expressed α-syn. We demonstrate that the LRRK2 G2019S mutation increases formation of α-syn aggregates in human iNs derived from fPD patients (Fig. 7). Conversely, loss of LRRK2 decreased α-syn aggregation in human iNs. Ultimately, our data posit reduction of LRRK2 as a potential therapeutic approach to mitigate α-syn aggregation and promote neuronal integrity. Clinical trials of LRRK2 kinase inhibitors are currently underway [3, 22]. Interestingly, increased LRRK2 kinase activity has recently been reported in sporadic PD [17]. Our results showing that LRRK2 knockout decreases aggregation of α-syn raises the possibility that these inhibitors may be efficacious in sporadic PD and not only in PD cases caused by LRRK2 mutations.

**Fig. 7.**
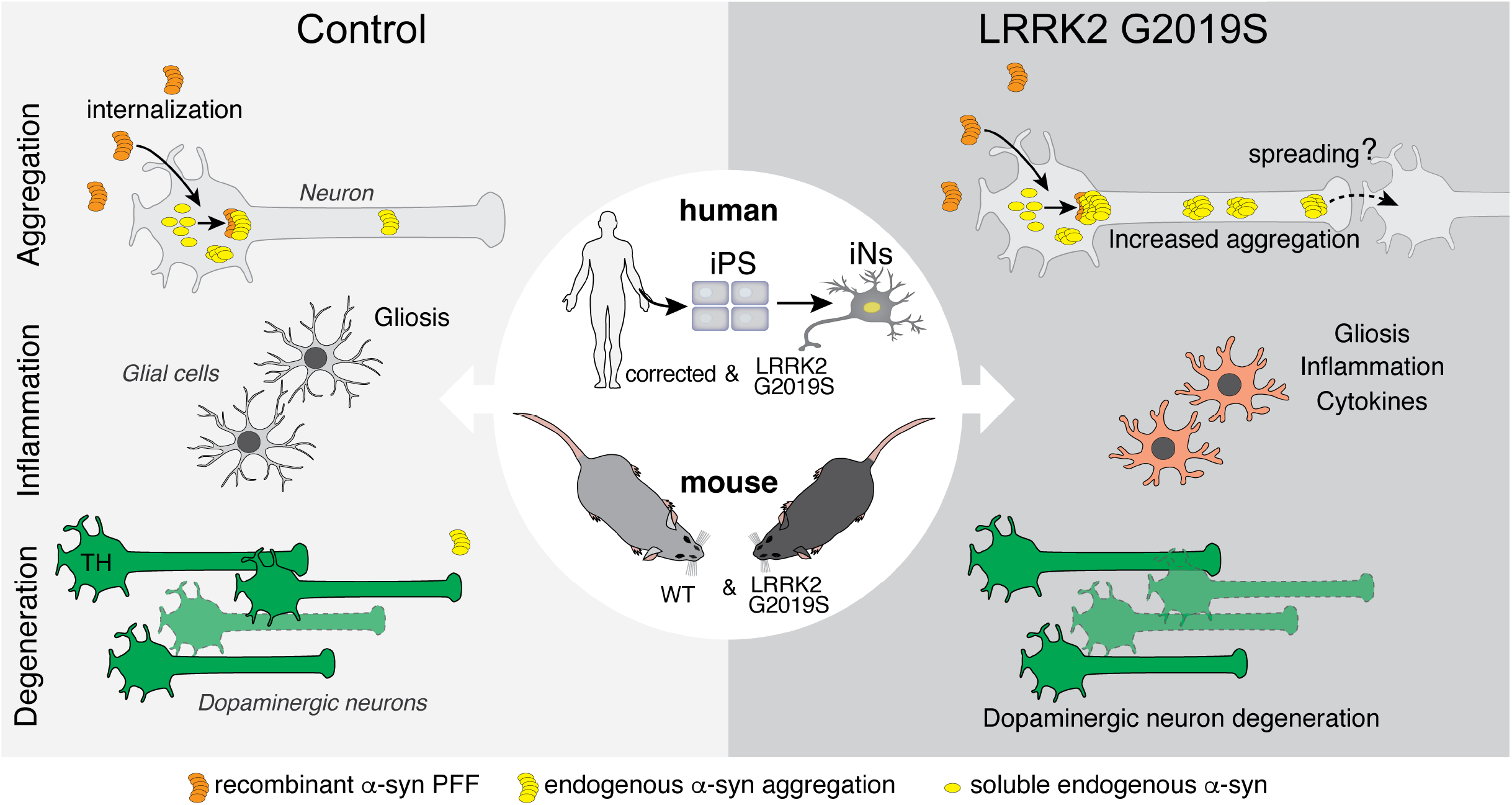
PD-linked mutations in LRRK2 exacerbate aggregation, glial activation and neuronal survival in mouse and human models of PFF-transmitted α-syn pathology. In primary neuron cultures and human iPS-derived induced neurons (iNs), PFFs are rapidly internalized and trigger the recruitment of soluble α-syn into cytoplasmic aggregates. The LRRK2 G2019S mutation is one of the most common genetic causes of Parkinson’s disease. Human and mouse neurons, derived from patient-derived iPS cells or LRRK2 G2019S BAC transgenic mice, develop normally *in vitro*. While internalization of PFFs is unaltered, LRRK2 G2019S neurons display increased accumulation of aggregated endogenous α-syn upon PFF treatment. Further, following intrastriatal PFF delivery into the brains, LRRK2 G2019S mice display an altered neuroinflammatory response with increased expression of inflammatory markers and cytokines and accelerated degeneration of dopaminergic neurons.

The PD risk gene-specific imaging-based screen described in our study enabled us to assess α-syn aggregation independent of toxicity and survival of primary neurons. Though focused in scope, our targeted screen successfully identified Gba and Lrrk2 as potential modifiers of PFF transmission and aggregation of endogenous α-syn. Interestingly, genes associated with juvenile forms of PD, such as Pink1 and Park2, which are not robustly associated with Lewy body pathology, exhibited no effects in this assay. Given the specificity of this imaging-based screen, its application could be expanded to identify additional modifiers or small molecule inhibitors of α-syn transmission and aggregation.

There are strong experimental and epidemiological evidence linking α-syn aggregation directly to endogenous levels of α-syn expression [40, 42]. In particular, genetic duplications and triplications of the *SNCA* locus are a genetic cause of fPD with exacerbated α-syn aggregation pathology and earlier disease onset [12, 54]. Through our screen, we demonstrated that knock down of Gba increased α-syn level in mouse primary neurons, possibly accounting for the observed increase in α-syn aggregation in response to PFF treatment. Consistent with our findings, interactions between Gba and α-syn levels have previously been reported in cell lines and primary neurons [2, 29, 44]. Moreover, GBA mutations are one of the most common genetic risk factors for PD in humans [68].

A growing number of animal models have been developed to further our understanding of pathogenesis of PD-associated LRRK2 mutations and α-syn aggregation. However, strong neuronal overexpression of both LRRK2 and human α-syn have yielded conflicting reports [14, 15, 32, 38]. It is unclear to what extent neuronal overexpression reflects the human pathology, given the ubiquitous expression of LRRK2 in the CNS in both neuronal and glial cells [65]. Additionally, PD is not commonly associated with elevated expression of α-syn, and overexpression of human α-syn can lead to toxic effects in eukaryotic cells independent of the formation of α-syn aggregates [13]. Recently, two studies investigated the effect of LRRK2 mutations using PFF-based *in vitro* models, both observing an increase in α-syn aggregation in primary neurons using the same transgenic mouse LRRK2 BAC line [30, 58]. Surprisingly, they report conflicting observations regarding the level of endogenously expressed α-syn protein level and the efficiency of LRRK2 inhibition in their mutant LRRK2 neurons. Although we initially identified the interaction between Lrrk2 and α-syn transmission in our targeted screen independent of the G2019S mutation, using a complementary human LRRK2 G2019S BAC-transgenic mouse strain we observe an increase in α-syn aggregation in primary neurons, in agreement with Volpicelli *et al*. and Henderson *et al* [30, 58]. We further extended our studies *in vivo*, delivering PFFs into the dorsal striatum of adult LRRK2 G2019S transgenic mice and observed accelerated α-syn aggregation and degeneration of TH neurons in our 6 month longitudinal study. Additionally, we observed an increase in the number of dopaminergic neurons with α-syn inclusions. Our findings are bolstered by a recent study in which PFFs were directly injected into the SNpc of rats, although this study does not account for transmission and transport of α-syn from a remote injection site to the site of aggregation [58]. Similarly, using a complementary AVV-based overexpression approach, α-syn burden and TH loss was recently shown to be exacerbated in an aging dependent manner in LRRK2 mutant mice [46]. Interestingly, while the distribution of α-syn pathology was surprisingly similar between LRRK2 and WT mice at 6 months post injection, we observed differing neuroinflammatory and glial responses in LRRK2 mutant mice at the same time point. Based on the growing knowledge of neuron-glia interaction in homeostasis, aging and neurodegenerative diseases, it is possible that glial cells could, in part, contribute to the accelerated phenotypes and impaired behavior observed in LRRK2 G2019S transgenic mice. Indeed, glial cells have recently been shown to contribute to α-syn pathology in a PFF-based mouse model [63], and several studies have proposed potential immune-related roles for LRRK2. For example, altered inflammatory and degenerative phenotypes have been reported in LRRK2-mutant rodents following LPS challenges [16, 45]. Taken together, these findings warrant further investigations into the relationship between LRRK2 mutations, accelerated α-syn pathology, and glial cells in models of PD.

Because of the success of PFF-based *in vitro* and *in vivo* mouse models and the increasing availability of human PD-iPS lines, we set out to develop a novel, scalable and robust PFF-based *in vitro* model for human neurons. We used a rapid induced neuron paradigm, allowing for the rapid differentiation of iPS cells into iNs, which developed α-syn aggregates in response to PFF treatment. As primarily a synaptic protein, α-syn is upregulated with increased maturation and synaptogenesis during differentiation. Importantly, we observed a significant increase in α-syn level and aggregation within a few weeks of differentiation without the need for α-syn overexpression. An intriguing distinction pertaining to PD between humans and rodents is the difference in the amino acid sequence of α-syn. The A53T mutation, associated with familial PD in humans, is the native amino acid sequence in mice and rats, possibly accounting for accelerated aggregation in rodent neurons [50]. Further, familial PD is associated with an earlier onset and more severe aggregation phenotype in subjects with the A53T mutation [23]. Given disparities between mouse and human models of PD at a fundamental structural level with respect to α-syn, it is critical to translate findings in animal models to a human disease context. To that end, various iPS and fibroblast-based protocols have recently been developed, potentially allowing for the generation of different neuronal subtypes that include those associated with individual degenerative diseases [28, 57]. Furthermore, the relative ease of use, the homogeneity of the neuron cultures and the ability to scale up and generate sufficiently high numbers of cells make iNs strong candidates for novel human neuron-based platforms to interrogate underlying mechanisms of disease progression and identification of potential therapeutic targets. Indeed, we demonstrated that loss of LRRK2 decreased α-syn aggregation in PFF-treated fPD patient-derived iNs, complementing results from *in vivo* antisense oligonucleotide (ASO) approaches in mice [67], and lending credence to the suitability of LRRK2 as a potential drug target.

## Acknowledgements

This work was supported by the National Institutes of Health grant R35NS097263 (A.D.G), Stanford BioX (A.D.G), research funding from Novartis Institutes of Biomedical Research (A.D.G), Stanford Graduate Fellowship (GB), HHMI fellowship (GB). RM and LB lab was supported by Grants from the EC Joint Programme on Neurodegenerative Diseases (JPND-SYNACTION-ANR-15-JPWG-0012-03), the Fondation pour la Recherche Médicale (Contract DEQ 20160334896) and the Innovative Medicine Initiative 2 grant agreement No 116060 (IMPRiND, www.imprind.org) supported by the European Union’s Horizon 2020 research and innovation program and EFPIA. The generation of the patient-specific ZFN-modified LRRK2 lines were supported by the California Institute of Regenerative Medicine (RT2-0196, BS). We thank Dr. Marius Wernig and Dr Daniel Haag for providing the TetOn and Ngn2 plasmids and advice with the induced neuron platform, Dr. Herschel and Dr. Suzanne Pfeffer for advice on LRRK2 expression and activity assays and Dr. Laura Volpicelli-Daley for advice on pSer129 immunostainings and immunoblotting. We thank A. Olsen and the Stanford Neuroscience Microscopy Service, supported by a grant from NIH (NS069375) and J. Mulholland from the CSIF for help with microscopy and image acquisition, Y. Zuber for mouse husbandry advice and support, the High Throughput Bio-Science center for providing access to imaging and analysis tools.

## Ethical approval

All applicable international, national, and/or institutional guidelines for the care and use of animals were followed. All procedures involving animals were in accordance with the ethical standards set by the Administrative Panel on Laboratory Animal Care of Stanford University (APLAC). This article does not contain any study with human participants performed by any of the authors.

