## Supplemental Data for "LRRK2 modifies α-syn pathology and spread in mouse models and human neurons"

**Supplementary Material**

### Figure Legends – Supplementary Figures

**Figure S1: Recombinant α-syn PFFs induce aggregation of endogenous α-syn in primary neurons *in vitro.***

**a** Schematic of lentiviral or PFF-based α-syn aggregation assay. Primary cortical neurons from E17 mouse embryos were infected with α-syn expressing lentivirus or treated with 2.6ug/ml sonicated mouse α-syn PFFs at 10 days in vitro (DIV). Neuronal survival and pSer129-α-syn aggregation was measured 1, 3, 6, 10 days post PFF/lentiviral treatment. **b,c** Neuronal survival was measured using luminescence-based LDH activity assays **(b)** and CaspaseGlo DEVDase assays **(c)** 10 days after lentiviral infection or PFF addition to the culture media. **d,e** Representative images **(d)** and quantification **(e)** of pSer129-α-syn aggregation (green) 3-10 days post PFF treatment. MAP2 (magenta) was used as a marker for neuronal processes (n=8 wells/time point; 12 images/well). Data expressed as mean + SEM; $*p < 0.05$; $**p < 0.01$; $***p < 0.001$; compared by one-way ANOVA with a Tukey's post-test for multiple comparisons.

**Figure S2: Characterization and size measurements of sonicated human and mouse α-syn preformed fibrils.**

**a** Representative electron micrographs of sonicated mouse α-syn preformed fibrils (PFFs). The scale bar, 100nm. **b** Quantification and distribution of fragment lengths, in 10nm bins. Average fragment length of the mouse PFFs is 50.39nm. **c** Representative electron micrographs of sonicated human α-syn PFFs. **d** Quantification and distribution of fragment length, in 10nm bins. The average fragment length of the human PFFs is 50.4nm.

**Figure S3: Targeted Lentiviral knockdown of Parkinson's disease risk genes in primary neurons.**

**a-f** Lentivirus-mediated knock-down efficiency was measured using quantitative RT-qPCR analysis. Neurons were infected on 1 day in vitro (DIV) and mRNA was isolated on 10 DIV. 4-5 different shRNA constructs were generated per Parkinson's disease risk gene. Representative knock down efficiency of shRNAs targeting Lrrk2 **(a)**, Gba **(b)**, Snca **(c)**, non-targeting scramble control **(d)**, Pink1 **(e)** and Park2 **(f)**, normalized to Scramble control (n=3-4 wells/shRNA). Data expressed as mean + SEM; $*p < 0.05$; $**p < 0.01$;$***p<0.001$ compared by one-way ANOVA with a Tukey's post-test for multiple comparisons.

**Figure S4: Characterization and manipulation of Gba and Lrrk2 in primary neurons *in vitro.***

**a,b** Representative Western blot **(a)** and quantifications **(b)** of lysates from primary neurons infected with Gba and non-targeting scramble shRNAs (SCR), collected 9 days post infection. Blots were probed with anti-α-syn, anti-ActB, anti-Lrrk2 and anti-Gba antibodies. ActB was used as a loading control and the data normalized to the scramble condition (n=4/group). Data expressed as mean + SD. **c** Schematic of experimental design: Primary neurons were isolated from CAG-Cas9 transgenic E17 embryos, infected with Lrrk2 and non-targeting (NT) gRNAs on 1 DIV and treated with sonicated α-syn PFF at 10DIV. **d** α-syn aggregation and viability was measured using pSer129 and NeuN immunolabeling 10 days post fibril treatment (n=10 wells/group). **e** Relative α-syn (Snca) mRNA expression level measured in NT and Lrrkt gRNA infected neurons at 10DIV using quantitative RT-PCR analysis. Data was normalized to the NT condition (n=6 wells/group) **f** Representative Sanger sequencing chromatogram of the G to A nucleotide mutation in exon 41 of the human LRRK2 in the BAC transgenic mouse line. Mutation marked in red. **g,h** Relative mRNA expression level of the synaptic markers Synaptophysin **(g)** and Snap25 **(h)** in WT compared to LRRK2 G2019S transgenic primary neurons at 10 DIV (n=6 wells/group). **i,j** Neuronal survival of α-syn PFF and Vehicle (Veh) treated WT compared to LRRK2-G2019S primary neurons was measured using LDH **(i)** and CaspaseGlo **(j)** luminescence-based activity assays (n=12 wells/group). **k** Quantification of internalization in G2019S LRRK2 and WT primary neuron cultures after 1-24 hours of incubation with fluorescently labeled sonicated α-syn PFFs (n=4 wells, 112-145 cells/well). Data expressed as mean + SEM; $*p < 0.05$; compared by unpaired Student's t-test and one-way ANOVA with a Tukey's post-test for multiple comparisons and two-way ANOVA with Bonferroni post hoc correction.

**Figure S5: Recombinant α-syn PFF delivery induce aggregation of endogenous α-syn in the brain of adult wildtype mice.**
**a** Experimental design: adult wildtype (WT) mice were stereotaxically injected with recombinant mouse α-syn PFFs or vehicle control (Veh) into the dorsal striatum, followed by behavioral testing and histological analysis up to 6 months post injection (PI). **b** Representative images of pSer129-postive (green) and p62-positive (red) α-syn aggregates and DAPI (blue) in the dorsal striatum 6 months post PFF injection. **c** Schematic of coronal brain section with motor cortex and representative confocal image of Thioflavin S staining (ThioS, green) and pSer129 immunolabeling (red). Scale bar, 100μm. **d** Schematic of coronal brain section with Striatum and representative confocal image of Thioflavin S staining and pSer129 immunolabeling. Scale bar, 50μm. **e** Schematic of coronal brain section with Amygdala and representative confocal image of Thioflavin S staining and pSer129 immunolabeling. Scale bar, 100μm. **f** Schematic of coronal brain section with Substantia nigra pars compacta (SNpc) and representative confocal image of Thioflavin S staining and pSer129 immunolabeling. Scale bar, 100μm.

**Figure S6: Intrastriatal delivery of PFF triggers α-syn pathology, degeneration of dopaminergic neurons and changes in open field behavior assays.**
**a** Schematic of a coronal mouse brain section with the substantia nigra pars compacta highlighted (gray). **b** Representative images of pSer129-postive aggregates (green) in TH-positive (red) dopaminergic neurons in the SNpc 2 months post PFF injection (PI). **c** Quantification of percentage of TH-positive neurons with pSer129-positive inclusions (n=8-12 animals/group). **e** Representative images of TH-positive (red) dopaminergic neurons in SNpc ipsilateral and contralateral to the injection site at 2 and 6 months post injection with PFFs. **e,f** Quantification of TH-positive neurons in in the SNpc of PFF **(e)** and vehicle control **(f)** injected animals (n=6-12 animals/group, 8-10 sections/animal). Black bars indicated the number of TH-positive neurons ipsilateral to the injection site, gray bars contralateral to the injection site. Data expressed as mean + SD. **g-i** Behavioral testing of PFF and vehicle control injected mice up to 6 months post injection. **g** Rotarod test measured the latency to fall from an accelerating rotating cylinder (n=10-12 animals/group). **h** Open field measures anxiety using the time spent in the center and the overall distance travelled (n=10-12 animals/group). Data expressed as mean + SEM; $*p < 0.05$; compared by Student's t-test, one-way ANOVA with a Tukey's post-test for multiple comparisons or two-way ANOVA with Bonferroni post hoc correction.

**Figure S7: Characterization of G2019S LRRK2 BAC transgenic mice.**
**a** Schematic illustrating the two mouse genotypes characterized: LRRK2 G2019S BAC transgenic mice and WT littermate controls. **b** Representative Western blot of striatal brain lysates from LRRK2 G2019S-BAC transgenic (G2019S or GS) mice and WT littermate controls were probed

with anti-human LRRK2, anti-α-syn and anti-ActB antibodies. ActB was used as loading control. **c** Quantification α-syn protein level in G2019S-LRRK2 BAC transgenic mice and WT littermate controls (n=4 animals/group). **d** qRT-PCR expression analysis of α-syn mRNA level in 2 months old G2019S-LRRK2 BAC transgenic mice and WT littermate controls (n=4 animals/group). **e** Rotarod behavioral assay measuring the latency to fall from an accelerating rotating cylinder for 2, 6, 12 months old G2019S-LRRK2 BAC transgenic mice and WT littermate controls (n=12-15/group). **f** Body weight measurements of 2 to 12 months old G2019S-LRRK2 BAC transgenic mice and WT littermate controls (n=20-25/group). Data expressed as mean + SEM or mean + SD; n.s.,$p > 0.05$; compared by Student's t-test, one-way ANOVA with a Tukey's post-test for multiple comparisons and two-way ANOVA with Bonferroni post hoc correction.

**Figure S8: The α-syn aggregation pathology is exacerbated in LRRK2 G2019S BAC transgenic mice injected with α-syn preformed fibrils.**
**a** Schematic representation of analyzed brain area, representative images and quantification of α-syn pSer129 labeling in the dorsal Striatum at 1, 3 and 6 months post injection with α-syn PFFs or vehicle (Veh) control into LRRK2 G2019S mutant (G2019S or GS) and wildtype (WT) littermate control mice (n=6-12 animals/group). Scale bar, 200µm. **b** Schematic representation of brain area, representative images and quantification of α-syn pSer129 labeling in the motor cortex at 1, 3 and 6 months post injection with α-syn PFFs or vehicle control into LRRK2 G2019S and WT mice. (n=6-12 animals/group). Scale bar, 250µm. **c** Schematic representation of brain area, representative images and quantification of α-syn pSer129 labeling in the basolateral Amygdala at 1, 3 and 6 months post injection with α-syn PFFs or vehicle control into LRRK2 G2019S and WT mice. (n=6-12 animals/group). Scale bar, 200µm. Data expressed as mean + SEM; *$p < 0.05$; compared by Student's t-test, one-way ANOVA with a Tukey's post-test for multiple comparisons.

**Figure S9: Rapid differentiation and upregulation of synaptic and mature neuron markers using a Ngn2-based induced neuron approach.**
**a** Bright field images of undifferentiated iPS colonies and induced neurons (iNs) after two weeks of differentiation *in vitro*. Schematic and time-line of Ngn2-based differentiation protocol. iPS cells differentiate rapidly into iNs following Ngn2 expression and transition into neuronal differentiation media. **b,c** Representative images of iNs expressing neuronal markers B-III-tubulin **(b)** and MAP2 **(c)**. **d** qRT-PCR analysis of human endogenous α-syn (SNCA) mRNA expression in iPS control lines during 4 weeks of iN differentiation (n=4 iPS lines/time point). Data normalized to 0 weeks of differentiation. **e** Representative Western blot of lysates form iNs up to 4 week of differentiation probed with anti-α-Syn and anti-ACTB antibodies. ACTB was used as a loading control. **f-i** RT-qPCR expression analysis of mature neuron markers MAP2 **(f)** MAPT **(g)** and synaptic markers SNAP25 **(h)** and GRIA1 **(i)** during 4 weeks of iN differentiation (n=4 iPS lines/time point). Data normalized to the 0 weeks of differentiation time point. Data expressed as mean + SEM.

**Figure S10: Survival analysis of PD-patient derived isogenic LRRK2 G2019S and knock-out iNs.**
**a** Representative Sanger sequencing chromatogram of the G to A nucleotide mutation in exon 1 of LRRK2 in G2019S LRRK2 mutant (top) and healthy control (bottom) iPS lines. **b,c** Neuronal survival of isogenic G2019S LRRK2 mutant (G2019S), corrected and LRRK2 knock-out (KO) iNs was measured after 4 weeks of differentiation using LDH **(b)** and CaspaseGlo DEVDase **(c)** luminescence-based activity assays (n=12 wells/line/time point). Data expressed as mean + SD; n.s.,$p>0.05$; compared by one-way ANOVA with a Tukey's post-test for multiple comparisons.

Bieri_Figure S1

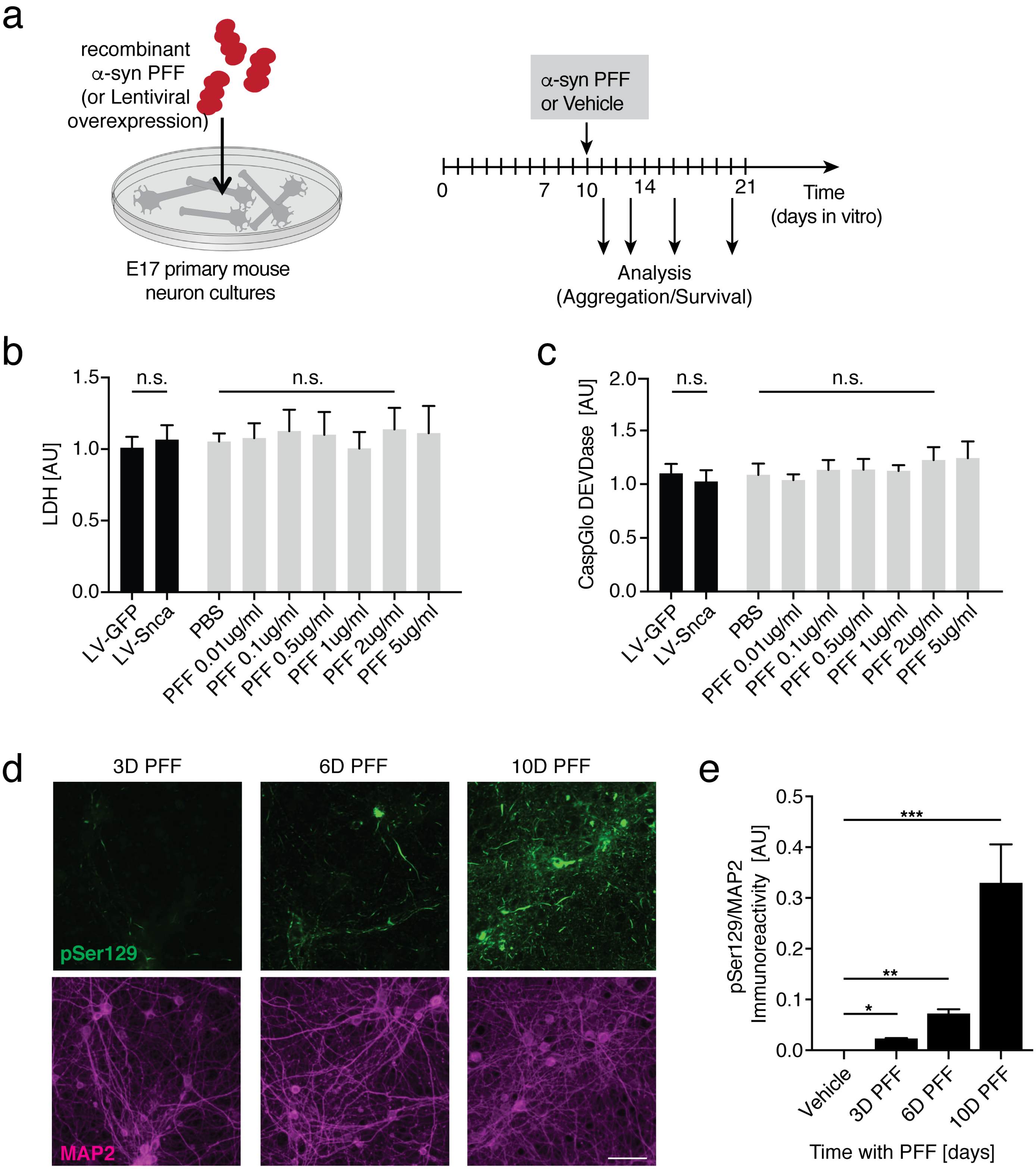

Bieri_Figure S2

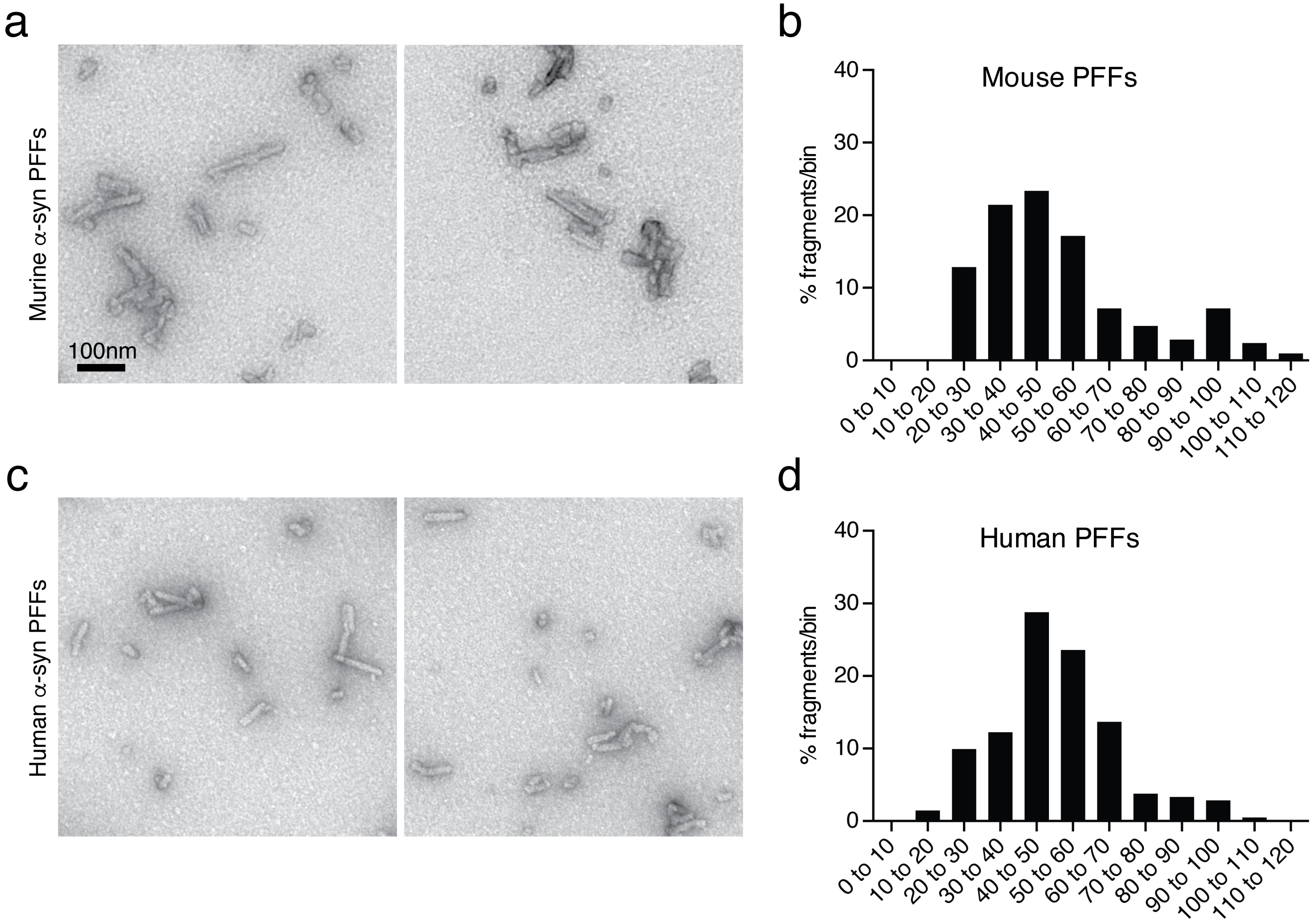

Bieri_Figure S3

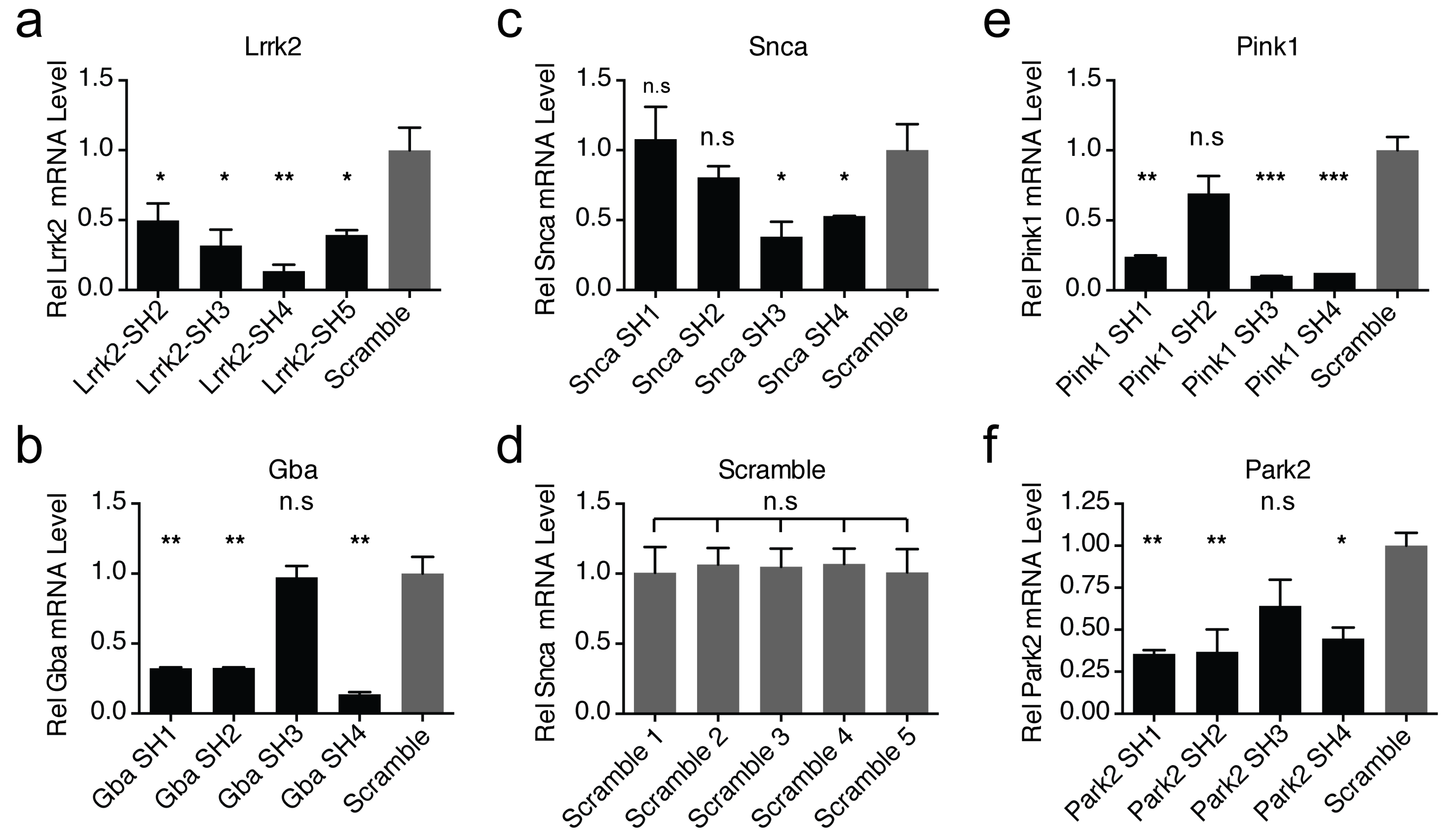

Bieri_Figure S4

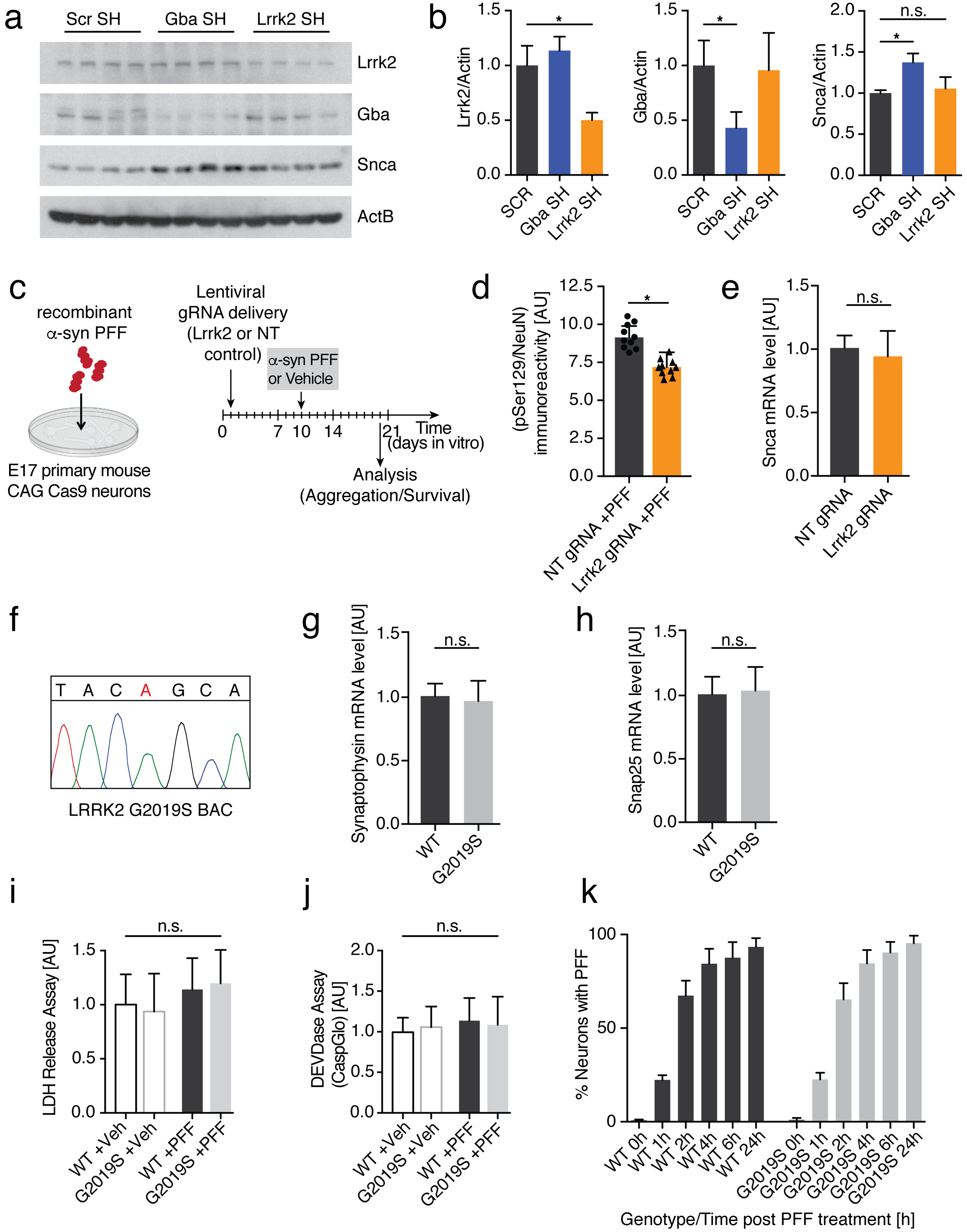

Bieri_Figure S5

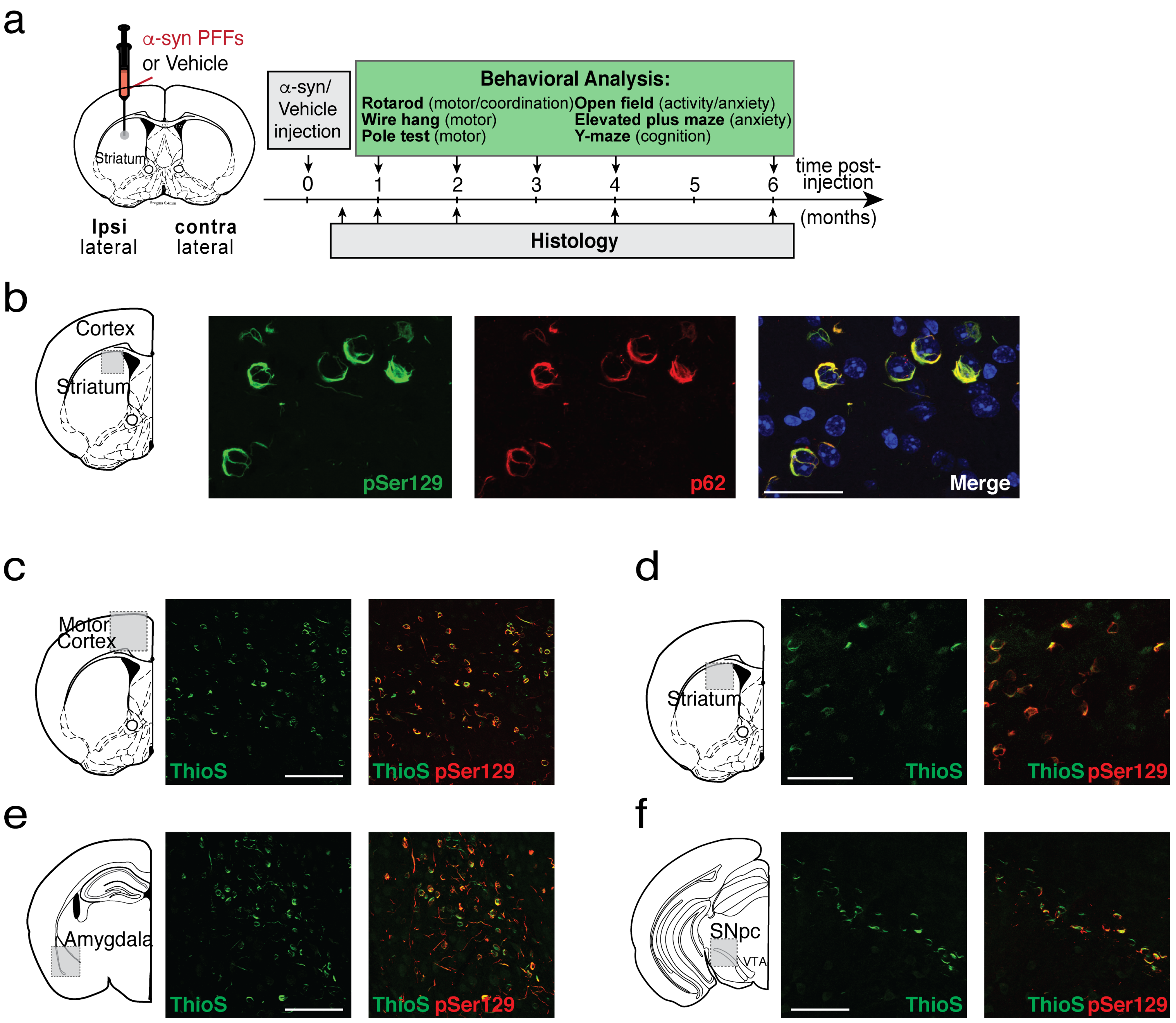

Bieri_Figure S6

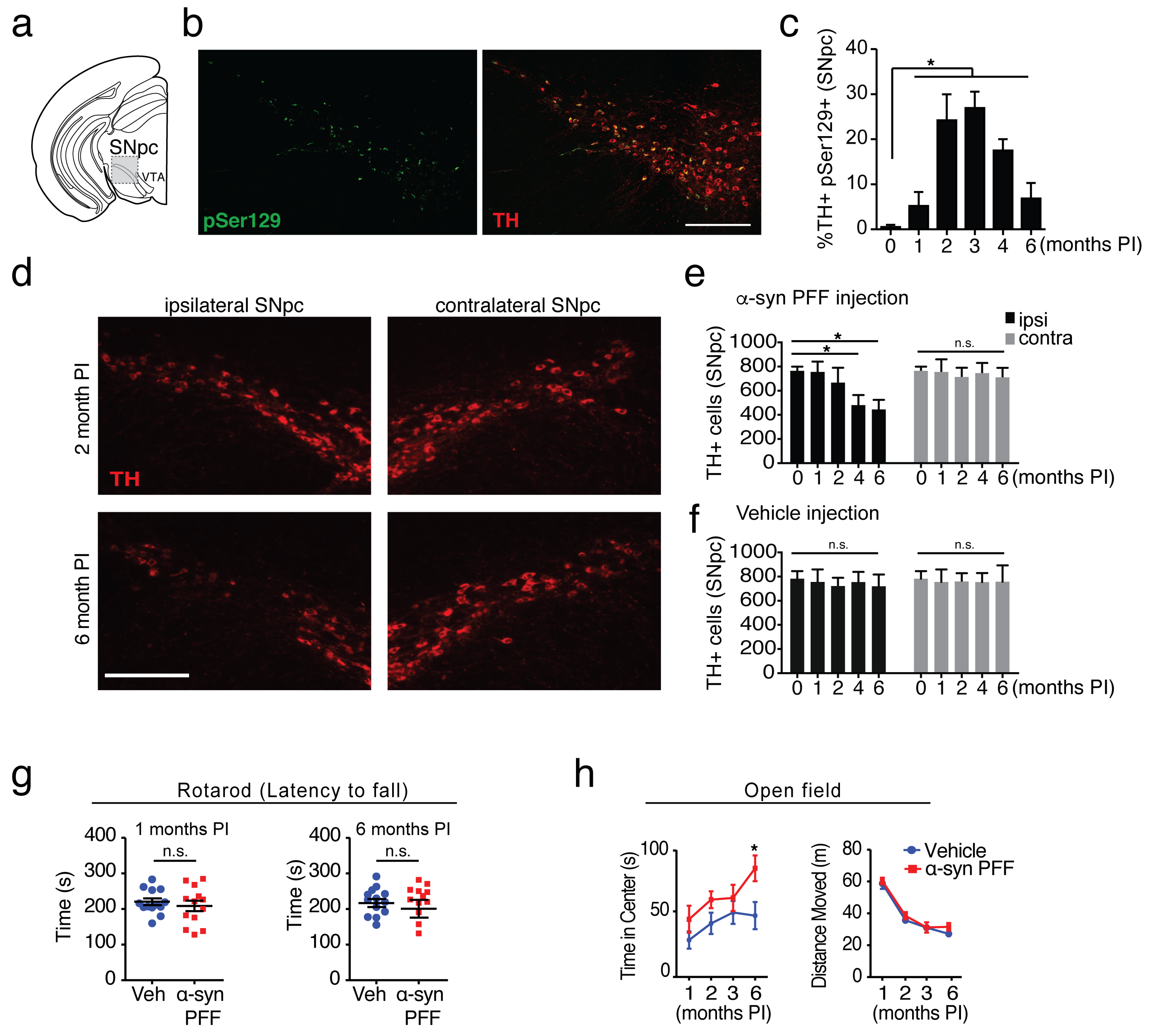

Bieri_Figure S7

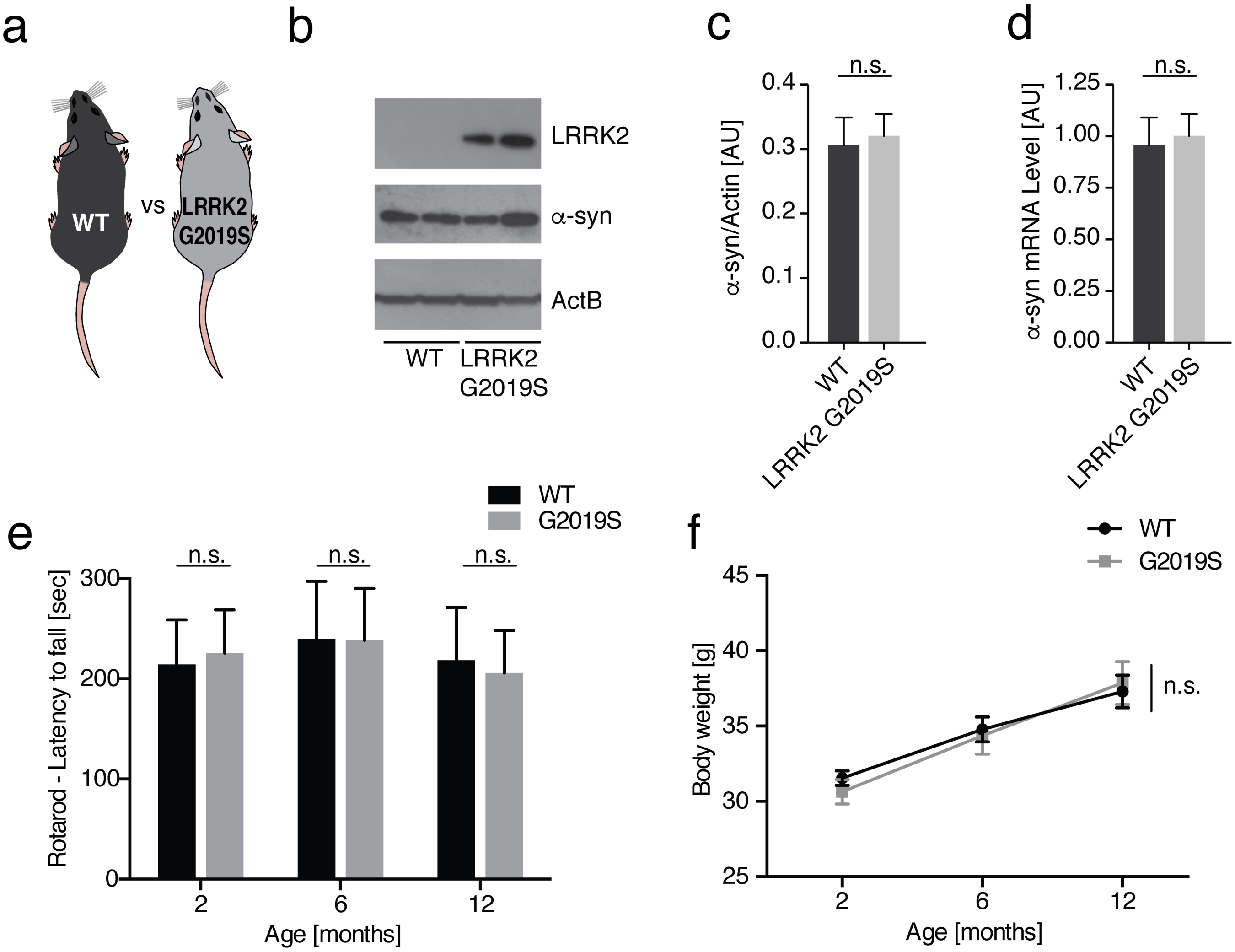

Bieri_Figure S8

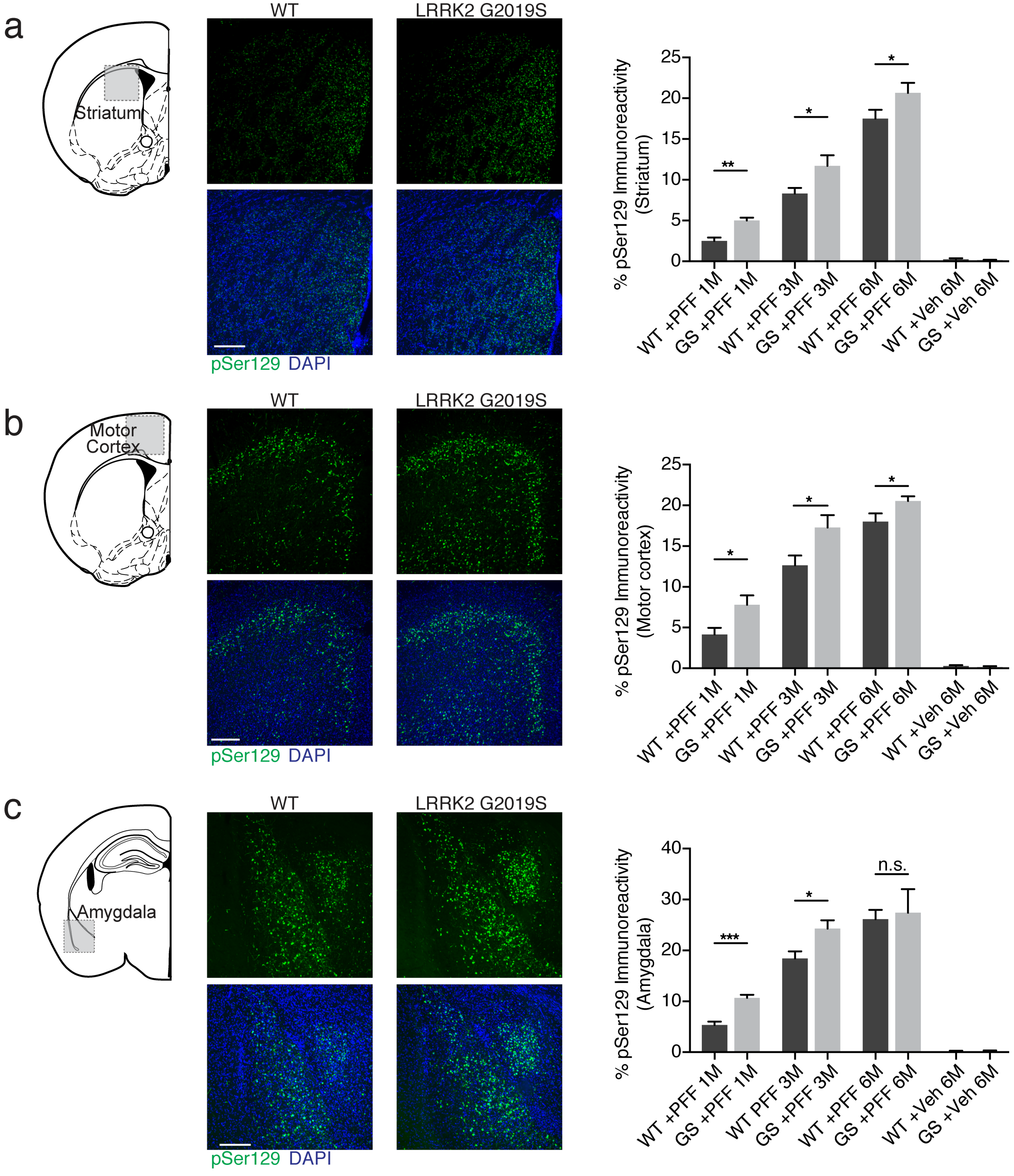

Bieri_Figure S9

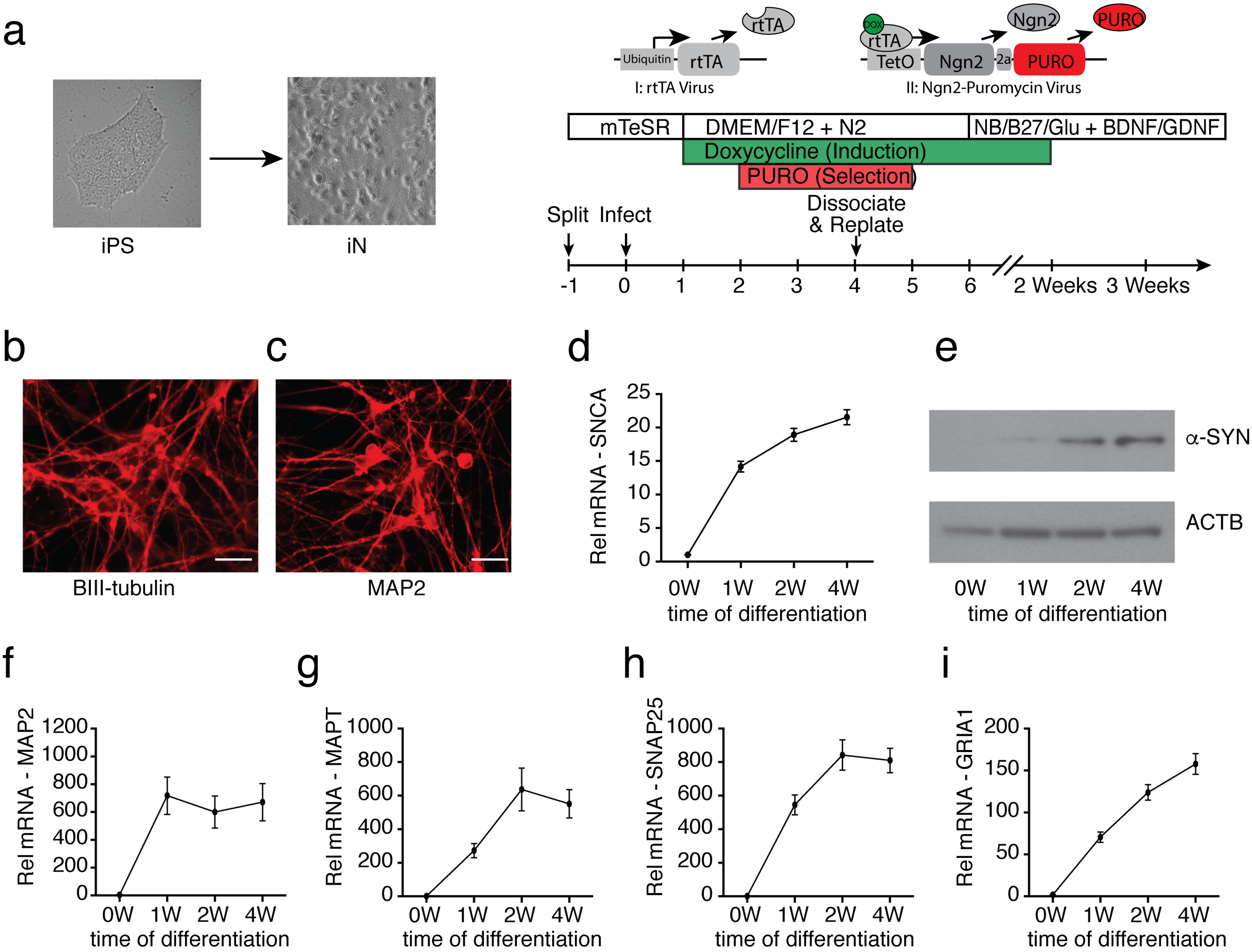

Bieri_Figure S10

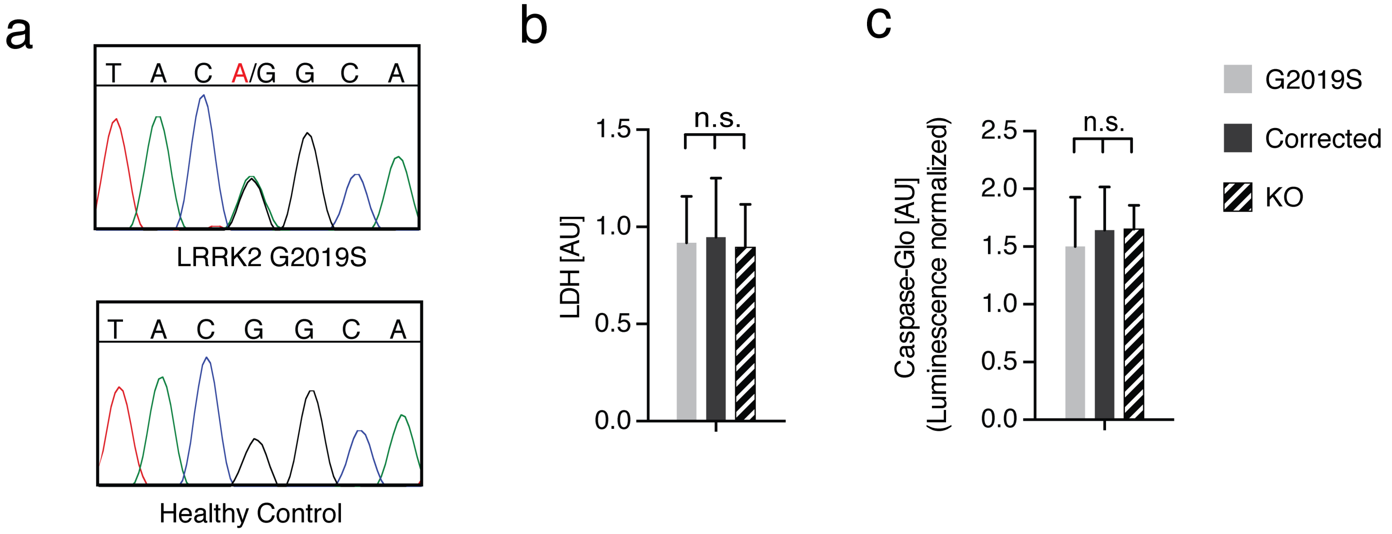
